# How medically important antimicrobials bind to the 30S ribosomal subunit in a bacterial pathogen

**DOI:** 10.64898/2025.12.24.696192

**Authors:** Swati R. Manjari, Caleb Mallery, Nilesh K. Banavali

## Abstract

Ribosomes translate the genetic code in mRNA to synthesize proteins in all living organisms. Decoding of mRNA occurs in the small subunit of the ribosome and is mediated by tRNA anticodons. Regions near the decoding center are a target for antibiotics, such as aminoglycosides and tetracyclines, where their presence results in errors in protein synthesis. More than two decades of high-resolution structural studies have shown how such medically important antimicrobials (MIAs) bind to the small subunit of the bacterial ribosome. Here, we comprehensively analyze these previously reported structures to help understand the variability with which MIAs bind to small subunits of bacterial ribosomes. We previously solved the hibernating 70S ribosome structure of the bacterial pathogen *Borrelia burgdorferi* (*Bbu*), the causative agent of Lyme disease, but there is no structure of any MIA bound to this ribosome reported. Our structural analysis makes it possible to use inexpensive computational methods to predict binding of these MIAs to the *Bbu* ribosomal 30S small subunit. For this, we used structural analogy, restrained energy minimization, and single-point binding free energy computations. We find the single-point binding free energy of the MIAs to be very sensitive to small conformational changes in the MIA and its environment. Incorporating this knowledge in structure-guided design could aid in development of narrow-spectrum MIAs targeting specific bacterial pathogens.

---

Ribosomes are essential and abundant molecular complexes in all living cells that translate a genetically message encoded in three base increments (codons) in mRNA to synthesize the corresponding proteins^1,2^. The mRNA codons are “decoded” by their complementary anticodons in aminoacyl tRNAs (aa-tRNAs) at a site called the decoding center (DC) in the small subunit (30S subunit in bacteria)^3^. The mechanism of action of many antimicrobials involves interfering with bacterial translation by binding to their ribosomes^4^. Many ribosomal small subunit binding antimicrobials bind near the DC^5^, and alterations near their binding sites can cause antibacterial resistance^6,7^. Antibacterial resistance is an increasing threat that is anticipated to result in high mortality and economic burdens^8^ unless it is addressed. In this regard, the World Health Organization (WHO) publishes a list of <u>m</u>edically important <u>a</u>ntimicrobials (MIAs) needed to treat bacterial infections in humans and animals^9^.

A large majority of ribosomal small subunit binding MIAs consist of aminoglycosides or tetracycline derivatives^9^. MIA aminoglycosides that target bacterial ribosomal small subunits are in wide-spread clinical use^10^. They have amino modified sugars and are classified into two classes based on whether the aminocyclitol moiety 2-deoxy streptamine (2-DOS) is present or not^11^. The antibiotics containing 2-DOS bind to the 16S rRNA helix 44 (h44) and stabilize flipping of the rRNA nucleotides, A1492 and A1493, in *Escherichia coli (Eco)* numbering^12^. These two highly conserved bases interact with the minor groove of the RNA duplex between the tRNA anticodon and the mRNA codon to probe cognate tRNAs during decoding^3^. These aminoglycosides affect translation by stabilizing non-cognate tRNA interaction with the mRNA, inhibiting translocation, and reduce subunit splitting during ribosome recycling^13^. Aminoglycosides not containing the 2-DOS moiety, such as streptomycin, bind to a different site on the small subunit composed of rRNA helices h1, h18, h27 and h44, and the protein S12^14^. Streptomycin is also thought to stabilize non-cognate tRNA-mRNA duplexes, but by altering the conformation of the decoding site, and not by inducing A1492 and A1493 flipping^15^. Tetracyclines bind to a site near rRNA helix h31, where the anticodon stem loop of the A-site tRNA binds during translation^12^.

Recent advances in deep learning-based approaches have yielded rapid and high-accuracy prediction of macromolecular structures^16–18^. This accuracy suggests that the simplified energy functions^19–21^ used to model the molecules include sufficient detail for reliable structure prediction when combined with sequence covariation-derived restraints. Supplementing prior structural knowledge with structure prediction of individual components with unknown structure can yield accurate models for macromolecular complexes through integrative modeling^22^. The binding of individual inhibitors can be modeled, and the accuracy of the modeling can be validated through comparison to subsets of these inhibitors that have known bound structures^23^. Structures for small molecule binding to targets with no experimentally determined structures can also be predicted, and knowledge of the structural features of the environment where they bind can be used to alter them for greater specificity or efficacy^24,25^. These advances should therefore enable accurate prediction of the binding of any MIA to any ribosome.

In this work, we analyze the conformational variability in the MIAs and their binding sites in known MIA bound ribosomal small subunit structures. We apply this knowledge to predict MIA binding to the 30S ribosomal subunit from a pathogen species: *<u>B</u>orrelia <u>bu</u>rgdorferi* (*Bbu*). We previously solved the structures of the hibernating *Bbu* 70S ribosome and a *Bbu* 50S subunit^23^ with some ribosomal RNA (rRNA) disorder^26^, but there is no MIA-ribosome complex structure reported for *Bbu* yet. The details of how MIAs bind to the *Bbu* ribosome are important because one of them, doxycycline, is the first line treatment for Lyme disease, which occurs in about half a million people in the US annually due to *Bbu* infections caused by tick bites^27^.

Using our approach, we also predict *Bbu* 30S subunit-bound structures for 17 MIAs that have no previously reported structure bound to any ribosome. These *Bbu* 30S subunit-bound MIA structure predictions enable quick examination of binding poses of any MIA to any other ribosome through simple sequence-based structural alignment with the *Bbu* 16S rRNA. The computationally predicted single-point binding free energies of the MIAs are analyzed to understand the influence of the precise conformation of MIA and differences between their accommodation in the *Bbu* ribosome. This provides a detailed structural glimpse into how all these MIAs could bind to a particular bacterial ribosomal 30S subunit, which would otherwise require extensive efforts to characterize experimentally. This knowledge can aid in structure-based design of narrow-spectrum antibiotics that are selective towards ribosomes from specific bacterial pathogens.

## Methods

### Analysis of MIA-bound ribosomal structures

The list of MIAs from the WHO^9^ was manually examined to find MIAs that bound to ribosomal small subunits as part of their mechanism of action. Of these, the aminoglycoside MIAs are listed in Table 1 and the non-aminoglycoside MIAs are listed in Table 2. This division was made because ribosome binding of aminoglycosides, containing the 2-deoxy-streptamine (2-DOS) moiety, is known to be accompanied by a specific structural change - flipping out of two conserved adenine bases involved in cognate tRNA recognition^3^. To model binding of these 2-DOS aminoglycoside MIAs, it is therefore essential to first model this conformational change in the ribosomal small subunit. For aminoglycosides not containing the 2-DOS moiety (e.g. streptomycin) or other small subunit binding MIAs, there is no such significant conformational change to be accounted for. Structures containing all identified MIAs, deposited as of August 2024, were searched within the RSCB databank^28^. These structures were filtered using the presence of ribosomal small subunits to identify only ribosomal small subunit-bound structures whose 4-letter PDB IDs are listed in Tables 1 and 2. The details of these structures are listed in Tables 3 and 4. For crystal structures that had more than one ribosome in the asymmetric unit, each ribosome was treated as a separate structure for the purpose of the analysis.

**Table 1.** Medically Important Antibacterial (MIA) aminoglycosides analyzed.

| Name | Pubchem CID | 3D Structure | Ribosome-bound structure PDB IDs |
| --- | --- | --- | --- |
| amikacin | 37768 |  | 6YPU |
| apramycin | 3081545 |  | 4AQY, 7PJS, 7PJT, 7PJU, 7PJV, 7P JW, 7P JX, 7P JY, 7P JZ, 8CGD |
| arbakacin | 68682 |  | 8Q4F |
| astromicin | 5284517 |  |  |
| bekanamycin | 439318 | 6NP5 |  |
| dibekacin | 470999 |  | 8IFB, 8IFD |
| dihydrostreptomycin | 439369 |  |  |
| framycetin (neomycin) | 8378 |  | 4LF6, 4LFB, 4V52, 4V57, 4V9C, 4W29, 5A9Z, 5AA0 |
| gentamicin | 3467 | 6NP3 | 4LF4, 4LF9, 4V53, 4V55, 5OBM, 8CGU |
| isepamicin | 3037209 |  |  |
| kanamycin | 6032 |  | 8EV7 |
| micronomicin | 3037206 |  |  |
| netilmicin | 441306 |  |  |
| paromomycin | 165580 |  | 1FJG, 1IBK, 1IBL, 1N32, 1N33, 1VVJ, 1XMO, 1XNQ, 1XNR, 2UU9, 2UUA, 2UUB, 2UUC, 2UXB, 2UXC, 2UXD, 2VQE, 2VQF, 3T1H, 3T1Y, 4DR2, 4DR4, 4GKJ, 4GKK, 4JYA, 4KHP, 4L47, 4L71, 4LEL, 4LF7, 4LF8, 4LNT, 4LSK, 4LT8, 4P6F, 4P70, 4TUA, 4TUB, 4TUD, 4TUE, 4V51, 4V5C, 4V5D, 4V5G, 4V5L, 4V5Y, 4V8C, 4V8F, 4V8J, 4V97, 4WOI, 4WRA, 4WSD, 4WT8, 4X62, 4X64, 4X65, 4X66, 5BR8, 5EL6, 5EL7, 5IB7, 5IQR, 5NDV, 5TCU, 6AZ1, 6AZ3, 6CAO, 6DTI, 6MKN, 6MPF, 6MPI, 6OF6, 6OJ2, 6OPE, 6ORD, 7K00, 7OIZ, 8EIU, 8EMM, 8G6W, 8G7Q, 8G7R, 8OEQ, 8OH6, 8OI5 |
| plazomicin | 42613186 |  | 7LH5 |
| ribostamycin | 33042 |  |  |
| sisomicin | 36119 |  | 6CAP, 6CAR |
| streptomycin | 19649 |  | 1FJG, 4DR3, 4DR6, 4DR7, 4DUZ, 4DV1, 4DV3, 4DV5, 4DV7, 4JI1, 4JI3, 4JI8, 4NXN, 6RW4, 6RW5, 8QRK, 8QRM, 8QRN |
| tobramycin | 36294 |  | 4LFC |

The default 3D structure for each antibacterial is obtained from Pubchem. If a 3D structure did not exist there, it is obtained from the PDB (4-character PDB ID shown).

**Table 2.** MIA aminoglycoside ribosome-bound structure details.

| Name | PDB ID | Res. (Å) | Method | Org. | RNA chain | Structure notes | Cit. |
| --- | --- | --- | --- | --- | --- | --- | --- |
| amikacin (AKN) | 6YPU | 2.9 | Cryo-EM | *Aba* | 2,4 | 30S | 42 |
| apramycin (AM2) | 4AQY | 3.5 | X-ray | *Tth* | A | 30S | 43 |
|  | 7PJS | 2.4 | Cryo-EM | *Eco* | A | 70S | 44 |
|  | 7PJT | 6.0 | Cryo-EM | *Eco* | a | 70S | 44 |
|  | 7PJU | 9.5 | Cryo-EM | *Eco* | a | 70S | 44 |
|  | 7PJV | 3.1 | Cryo-EM | *Eco* | a | 70S | 44 |
|  | 7PJW | 4.0 | Cryo-EM | *Eco* | a | 70S | 44 |
|  | 7PJX | 6.5 | Cryo-EM | *Eco* | a | 70S | 44 |
|  | 7PJY | 3.1 | Cryo-EM | *Eco* | a | 70S | 44 |
|  | 7PJZ | 6.0 | Cryo-EM | *Eco* | a | 70S | 44 |
|  | 8CGD | 2.0 | Cryo-EM | *Eco* | a | 50S, **not used** | 45 |
| arbekacin (84G) | 8Q4F | 3.1 | Cryo-EM | *Eco* | A | 70S | np |
| astromicin |  |  |  |  |  |  |  |
| bekanamycin |  |  |  |  |  |  |  |
| dibekacin (84D) | 8IFB | 2.4 | Cryo-EM | *Eco* | A | 70S | 46 |
|  | 8IFD | 2.6 | Cryo-EM | *Hsa* | 2m | 80S, **not used** | 46 |
| dihydrostreptomycin |  |  |  |  |  |  |  |
| framycetin or<br>soframycin<br>(neomycin, NMY) | 4LF6 | 3.3 | X-ray | *Tth* | A | 30S | np |
|  | 4LFB | 3.0 | X-ray | *Tth* | A | 30S | np |
|  | 4V52 | 3.2 | X-ray | *Eco* | AA, CA | 70S | 47 |
|  | 4V57 | 3.5 | X-ray | *Eco* | AA, CA | 70S | 48 |
|  | 4V9C | 3.3 | X-ray | *Eco* | AA, CA | 70S | 49 |
|  | 4W29 | 3.8 | X-ray | *Tth* | CA, AA | 70S | 50 |
|  | 5A9Z | 4.7 | Cryo-EM | *Tth* | BA | 70S | 51 |
|  | 5AA0 | 5.0 | Cryo-EM | *Tth* | BA | 70S | 51 |
| gentamicin (LLL) | 4LF4 | 3.3 | X-ray | *Tth* | A | 30S | np |
|  | 4LF9 | 3.3 | X-ray | *Tth* | A | 30S | np |
|  | 4V53 | 3.5 | X-ray | *Eco* | AA, CA | 70S | 47 |
|  | 4V55 | 4.0 | X-ray | *Eco* | AA, CA | 70S | 47 |
|  | 5OBM | 3.4 | X-ray | *Sce* | 2, 6 | 80S, **not used** | 52 |
|  | 8CGU | 1.9 | Cryo-EM | *Eco* | A | 30S | 45 |
| isepamicin |  |  |  |  |  |  |  |
| kanamycin (KAN) | 8EV7 | 2.9 | X-ray | *Tth* | 1a, 2a | 70S | 53 |
| micronomicin |  |  |  |  |  |  |  |
| netilmicin |  |  |  |  |  |  |  |
| paromomycin (PAR) | 1FJG | 3.0 | X-ray | *Tth* | A | 30S | 14 |
|  | 1IBK | 3.3 | X-ray | *Tth* | A | 30S | 3 |
|  | 1IBL | 3.1 | X-ray | *Tth* | A | 30S | 3 |
|  | 1N32 | 3.0 | X-ray | *Tth* | A | 30S | 54 |
|  | 1N33 | 3.4 | X-ray | *Tth* | A | 30S | 54 |
|  | 1VVJ | 3.4 | X-ray | *Tth* | QA, XA | 70S | 55 |
|  | 1XMO | 3.3 | X-ray | *Tth* | A | 70S | 56 |
|  | 1XNQ | 3.1 | X-ray | *Tth* | A | 30S | 57 |
|  | 1XNR | 3.1 | X-ray | *Tth* | A | 30S | 57 |
|  | 2UU9 | 3.1 | X-ray | *Tth* | A | 30S | 58 |
|  | 2UUA | 2.9 | X-ray | *Tth* | A | 30S | 58 |
|  | 2UUB | 2.8 | X-ray | *Tth* | A | 30S | 58 |
|  | 2UUC | 3.1 | X-ray | *Tth* | A | 30S | 58 |
|  | 2UXB | 3.1 | X-ray | *Tth* | A | 30S | 59 |
|  | 2UXC | 2.9 | X-ray | *Tth* | A | 30S | 59 |
|  | 2UXD | 3.2 | X-ray | *Tth* | A | 30S | 59 |
|  | 2VQE | 2.5 | X-ray | *Tth* | A | 30S | 60 |
|  | 2VQF | 2.9 | X-ray | *Tth* | A | 30S | 60 |
|  | 3T1H | 3.1 | X-ray | *Tth* | A | 30S | 61 |
|  | 3T1Y | 2.8 | X-ray | *Tth* | A | 30S | 61 |
|  | 4DR2 | 3.3 | X-ray | *Tth* | A | 30S | 15 |
|  | 4DR4 | 4.0 | X-ray | *Tth* | A | 30S | 15 |
|  | 4GKJ | 3.3 | X-ray | *Tth* | A | 30S | 62 |
|  | 4GKK | 3.2 | X-ray | *Tth* | A | 30S | 62 |
|  | 4JYA | 3.1 | X-ray | *Tth* | A | 30S | 63 |
|  | 4KHP | 3.1 | X-ray | *Tth* | A | 30S | 64 |
|  | 4L47 | 3.2 | X-ray | *Tth* | QA, XA | 70S | 55 |
|  | 4L71 | 3.9 | X-ray | *Tth* | QA, XA | 70S | 55 |
|  | 4LEL | 3.9 | X-ray | *Tth* | QA, XA | 70S | 55 |
|  | 4LF7 | 3.2 | X-ray | *Tth* | A | 30S | np |
|  | 4LF8 | 3.2 | X-ray | *Tth* | A | 30S | np |
|  | 4LNT | 2.9 | X-ray | *Tth* | QA, XA | 70S | 55 |
|  | 4LSK | 3.5 | X-ray | *Tth* | QA, XA | 70S | 55 |
|  | 4LT8 | 3.4 | X-ray | *Tth* | QA, XA | 70S | 55 |
|  | 4P6F | 3.6 | X-ray | *Tth* | QA, XA | 70S | 65 |
|  | 4P70 | 3.7 | X-ray | *Tth* | QA, XA | 70S | 55 |
|  | 4TUA | 3.6 | X-ray | *Tth* | QA, XA | 70S | 66 |
|  | 4TUB | 3.6 | X-ray | *Tth* | QA, XA | 70S | 66 |
|  | 4TUD | 3.6 | X-ray | *Tth* | QA, XA | 70S | 66 |
|  | 4TUE | 3.5 | X-ray | *Tth* | QA, XA | 70S | 66 |
|  | 4V51 | 2.8 | X-ray | *Tth* | AA, CA | 70S | 67 |
|  | 4V5C | 3.3 | X-ray | *Tth* | AA, CA | 70S | 68 |
|  | 4V5D | 3.5 | X-ray | *Tth* | AA, CA | 70S | 68 |
|  | 4V5G | 3.6 | X-ray | *Tth* | AA, CA | 70S | 69 |
|  | 4V5L | 3.1 | X-ray | *Tth* | AA | 70S | 70 |
|  | 4V5Y | 4.5 | X-ray | *Tth* | AA, CA | 70S | 47 |
|  | 4V8C | 3.3 | X-ray | *Tth* | AA, BA | 70S | 71 |
|  | 4V8F | 3.3 | X-ray | *Tth* | CA, BA | 70S | 71 |
|  | 4V8J | 3.9 | X-ray | *Tth* | AA, CA | 70S | 72 |
|  | 4V97 | 3.5 | X-ray | *Tth* | AA, CA | 70S | 72 |
|  | 4WOI | 3.0 | X-ray | *Eco* | AA, DA | 70S | 73 |
|  | 4WRA | 3.1 | X-ray | *Tth* | 13, 1G | 70S | 74 |
|  | 4WSD | 3.0 | X-ray | *Tth* | 13, 1G | 70S | 74 |
|  | 4WT8 | 3.4 | X-ray | *Tth* | Ab, Bb | 70S | 75 |
|  | 4X62 | 3.5 | X-ray | *Tth* | A | 30S | 76 |
|  | 4X64 | 3.4 | X-ray | *Tth* | A | 30S | 76 |
|  | 4X65 | 3.4 | X-ray | *Tth* | A | 30S | 76 |
|  | 4X66 | 3.5 | X-ray | *Tth* | A | 30S | 76 |
|  | 5BR8 | 3.4 | X-ray | *Tth* | A | 30S | np |
|  | 5EL6 | 3.1 | X-ray | *Tth* | 13, 1G | 70S | 77 |
|  | 5EL7 | 3.2 | X-ray | *Tth* | 13, 1G | 70S | 77 |
|  | 5IB7 | 3.0 | X-ray | *Tth* | 13, 1G | 70S | 78 |
|  | 5IQR | 3.0 | Cryo-EM | *Eco* | 2 | 70S | 79 |
|  | 5NDV | 3.3 | X-ray | *Sce* | 2, 6 | 80S, **not used** | 52 |
|  | 5TCU | 3.9 | Cryo-EM | *Sau* | A | 70S | 80 |
|  | 6AZ1 | 2.7 | Cryo-EM | *Ldo* | 1 | 40S | 81 |
|  | 6AZ3 | 2.5 | Cryo-EM | *Ldo* | 1 | 60S | 81 |
|  | 6CAO | 3.5 | X-ray | *Tth* | A | 30S | 82 |
|  | 6DTI | 3.5 | X-ray | *Tth* | A | 30S | 83 |
|  | 6MKN | 3.5 | X-ray | *Tth* | A | 30S | 83 |
|  | 6MPF | 3.3 | X-ray | *Tth* | A | 30S | 83 |
|  | 6MPI | 3.3 | X-ray | *Tth* | A | 30S | 83 |
|  | 6OF6 | 3.2 | X-ray | *Tth* | QA, XA | 70S | 84 |
|  | 6OJ2 | 3.2 | X-ray | *Tth* | QA, XA | 70S | 84 |
|  | 6OPE | 3.1 | X-ray | *Tth* | QA, XA | 70S | 84 |
|  | 6ORD | 3.1 | X-ray | *Tth* | QA, XA | 70S | 84 |
|  | 7K00 | 2.0 | Cryo-EM | *Eco* | A | 70S | 85 |
|  | 7OIZ | 2.9 | Cryo-EM | *Eco* | A | 70S | 86 |
|  | 8EIU | 2.2 | Cryo-EM | *Eco* | A | 70S | 87 |
|  | 8EMM | 2.1 | Cryo-EM | *Eco* | A | 70S | 88 |
|  | 8G6W | 2.0 | Cryo-EM | *Eco* | A | 70S | 89 |
|  | 8G7Q | 3.1 | Cryo-EM | *Eco* | a | 70S | 90 |
|  | 8G7R | 2.8 | Cryo-EM | *Eco* | a | 70S | 90 |
|  | 8OEQ | 3.3 | X-ray | *Cal* | B, CM | 80S, **not used** | np |
|  | 8OH6 | 3.4 | X-ray | *Cal* | B, CM | 80S, **not used** | np |
|  | 8OI5 | 2.9 | X-ray | *Cal* | B, CM | 80S, **not used** | np |
| plazomicin (EDS) | 7LH5 | 3.3 | X-ray | *Tth* | AA, CA | 70S | 91 |
| ribostamycin |  |  |  |  |  |  |  |
| sisomicin (SIS) | 6CAP | 3.4 | X-ray | *Tth* | A | 30S | np |
|  | 6CAR | 3.4 | X-ray | *Tth* | A | 30S | 92 |
| streptomycin (SRY) | 1FJG | 3.0 | X-ray | *Tth* | A | 30S | 14 |
|  | 4DR3 | 3.4 | X-ray | *Tth* | A | 30S | 15 |
|  | 4DR6 | 3.3 | X-ray | *Tth* | A | 30S | 15 |
|  | 4DR7 | 3.8 | X-ray | *Tth* | A | 30S | 15 |
|  | 4DUZ | 3.7 | X-ray | *Tth* | A | 30S | np |
|  | 4DV1 | 3.9 | X-ray | *Tth* | A | 30S | np |
|  | 4DV3 | 3.6 | X-ray | *Tth* | A | 30S | np |
|  | 4DV5 | 3.7 | X-ray | *Tth* | A | 30S | np |
|  | 4DV7 | 3.3 | X-ray | *Tth* | A | 30S | np |
|  | 4JI1 | 3.1 | X-ray | *Tth* | A | 30S | 93 |
|  | 4JI3 | 3.4 | X-ray | *Tth* | A | 30S | 93 |
|  | 4JI8 | 3.7 | X-ray | *Tth* | A | 30S | 93 |
|  | 4NXN | 3.5 | X-ray | *Tth* | A | 30S | np |
|  | 6RW4 | 3.0 | Cryo-EM | *Hsa* | A | 28S | 94 |
|  | 6RW5 | 3.1 | Cryo-EM | *Hsa* | A | 28S | 94 |
|  | 8QRK | 6.7 | Cryo-EM | *Hsa* | A | 28S | 95 |
|  | 8QRM | 3.1 | Cryo-EM | *Hsa* | A | 28S | 95 |
|  | 8QRN | 3.0 | Cryo-EM | *Hsa* | A | 28S | 95 |
| tobramycin (TOY) | 4LFC | 3.6 | X-ray | *Tth* | A | 30S | np |

Res. = Resolution, Cit. = citation, np = no publication, RNA chain = chain(s) used to orient structures.

**Table 3.**
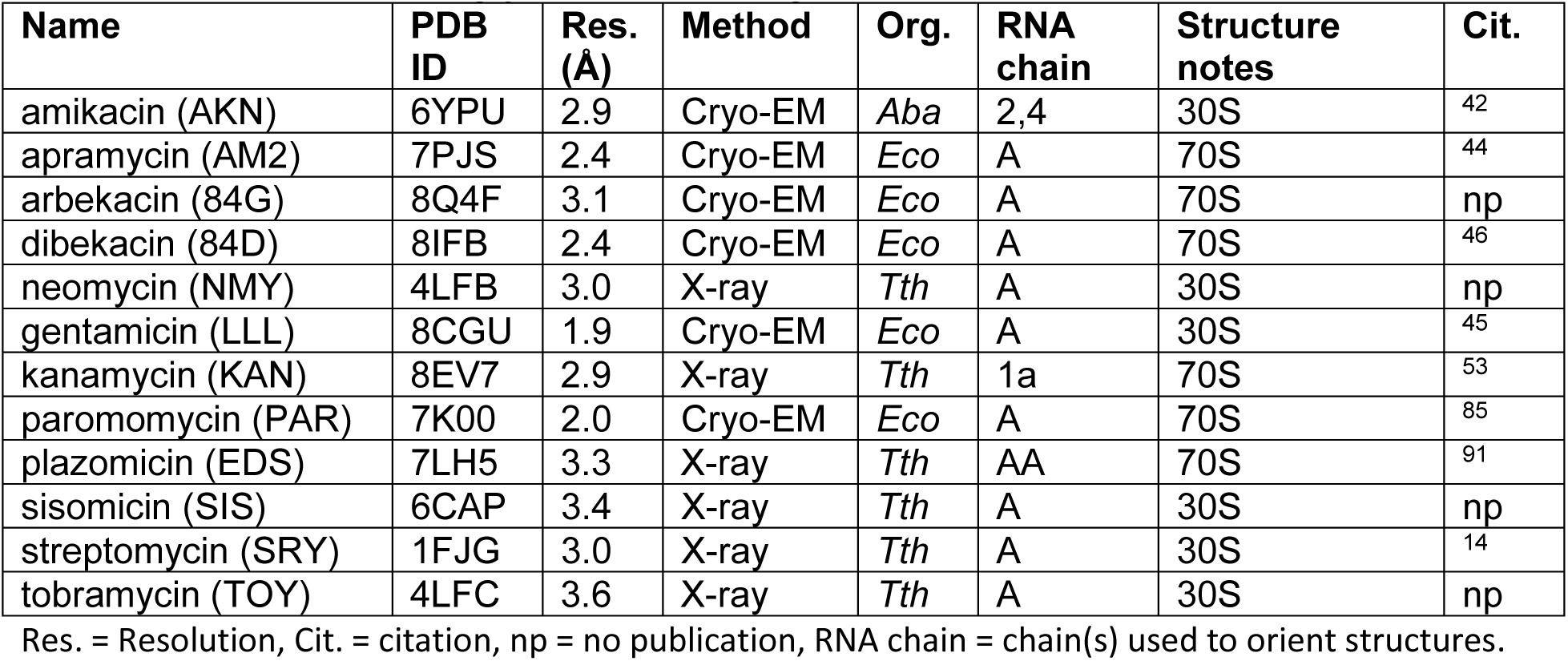
Template structures used for modeling *Bbu* 16S RNA structural changes to accommodate aminoglycoside binding.

| Name | PDB ID | Res. (Å) | Method | Org. | RNA chain | Structure notes | Cit. |
| --- | --- | --- | --- | --- | --- | --- | --- |
| amikacin (AKN) | 6YPU | 2.9 | Cryo-EM | *Aba* | 2,4 | 30S | <sup>42</sup> |
| apramycin (AM2) | 7PJS | 2.4 | Cryo-EM | *Eco* | A | 70S | <sup>44</sup> |
| arbekacin (84G) | 8Q4F | 3.1 | Cryo-EM | *Eco* | A | 70S | np |
| dibekacin (84D) | 8IFB | 2.4 | Cryo-EM | *Eco* | A | 70S | <sup>46</sup> |
| neomycin (NMY) | 4LFB | 3.0 | X-ray | *Tth* | A | 30S | np |
| gentamicin (LLL) | 8CGU | 1.9 | Cryo-EM | *Eco* | A | 30S | <sup>45</sup> |
| kanamycin (KAN) | 8EV7 | 2.9 | X-ray | *Tth* | 1a | 70S | <sup>53</sup> |
| paromomycin (PAR) | 7K00 | 2.0 | Cryo-EM | *Eco* | A | 70S | <sup>85</sup> |
| plazomicin (EDS) | 7LH5 | 3.3 | X-ray | *Tth* | AA | 70S | <sup>91</sup> |
| sisomicin (SIS) | 6CAP | 3.4 | X-ray | *Tth* | A | 30S | np |
| streptomycin (SRY) | 1FJG | 3.0 | X-ray | *Tth* | A | 30S | <sup>14</sup> |
| tobramycin (TOY) | 4LFC | 3.6 | X-ray | *Tth* | A | 30S | np |

Res. = Resolution, Cit. = citation, np = no publication, RNA chain = chain(s) used to orient structures.

**Table 4.** MIA non-aminoglycosides binding to ribosomal small subunits analyzed.

| Name | Pubchem CID | 3D Structure | Ribosome-bound structure PDB IDs |
| --- | --- | --- | --- |
| chlortetracycline | 54675777 |  |  |
| clomocycline | 54680675 |  |  |
| demeclocycline | 54680690 |  |  |
| doxycycline | 54671203 |  |  |
| eravacycline | 54726192 |  | 7M4U, 7M4V, 7M4W, 7M4X, 7M4Y, 7M4Z, 8CF8 |
| lymecycline | 54707177 |  |  |
| metacycline | 54675785 |  |  |
| minocycline | 54675783 |  |  |
| omadacycline | 54697325 |  | 8CA7 |
| oxytetracycline | 54675779 |  |  |
| pipacycline | 54686187 | * |  |
| rolitetracycline | 54682938 |  |  |
| sarecycline | 54681908 |  | 6XQD, 6XQE, 8CRX, 8CVM, 8CVO |
| spectinomycin | 15541 |  | 1FJG, 4V56, 4V57, 7N2V, 7P7Q, 7P7R, 7P7S, 7P7T, 7P7U, 7PIB, 8CA7, 8CAI, 8UVR |
| tetracycline | 54675776 |  | 1HNW, 1I97, 4V9A, 5J5B, 5J7L, 8CF1, 8CGJ |
| tigecycline | 54686904 | 4ABZ | 4V9B, 4YHH, 5J8A, 5J91, 6YSI, 6YTF, 8K2A, 8K2B, 8K2C, 8K2D, 8K82, 8XSX, 8XSY, 8XSZ, 8XT0, 8XT1, 8XT2, 8XT3, 8YOO |

The default 3D structure for each antibacterial is obtained from Pubchem but if a 3D structure does not exist there, it is either obtained from a PDB entry (4-character PDB ID shown) or generated using https://cactus.nci.nih.gov/translate from the SMILES in the Pubchem entry (each such instance indicated by \*).

### Modeling of the MIA-bound 30s subunit complexes for *Bbu*

The bound hibernation promotion factor (HPF) and E-site tRNA was removed from the hibernating 70S ribosome structure for *Bbu*^23^, and this was used as the base structure for the modeling of MIA-bound *Bbu* ribosome complexes. The first step for modeling MIA-bound ribosome complexes was aligning the entire 16S rRNA of each structure to the *Bbu* 16S rRNA using the matchmaker module of ChimeraX^29^ to find the approximate expected location of the MIA. In a second step, only the 16S rRNA residues within 15 Å of the MIA were chosen for structural alignment using matchmaker to provide a locally stringent alignment. If sequence identity of 16S rRNA residues with 15 Å was not sufficient for a good alignment (as judged against the full 16S rRNA alignment in the first step), the radius of the selection zone was increased to 20 Å. A total of 199 aminoglycoside and non-aminoglycoside structures (Tables 3 and 4) were aligned in this fashion to the *Bbu* ribosomal 30S subunit.

The MIAs examined bind in four sites in the 30S subunit, these accommodate: (a) the 2-DOS aminoglycoside analogs (Fig. 1A), (b) streptomycin and its analogs (Fig 1B), tetracycline and its analogs (Fig 1C), and spectinomycin and its analogs (Fig. 1D). Since there is variability in the precise conformation of the flipped adenine bases in different 2-DOS aminoglycoside bound ribosome structures (Fig. 1E), we used the highest resolution structures available for individual 2-DOS aminoglycosides as templates (Table 5, overlay shown in Fig. 1F) to generate corresponding homology models for the two adenine bases in their flipped conformations for *Bbu*. These template structures had the following MIAs bound to the ribosomes: amikacin, apramycin, arbekacin, dibekacin, neomycin, gentamicin, kanamycin, paromomycin, plazomicin, sisomicin, streptomycin, and tobramycin. The inclusion of the streptomycin-bound structure, even though streptomycin is not a 2-DOS aminoglycoside, is because its structure also has bound paromomycin that induces the two adenine base flipping, which happens in the vicinity of the bound streptomycin^14^. The 16S RNA residues 1485-1492 (*Bbu* numbering, sequence: GUG**<u>AA</u>**GUC with flipping adenines boldfaced and underlined) were homology modeled using the program ModeRNA^30^ using the corresponding residues in each of the template structures. This yielded 12 template structures of *Bbu* ribosomes with two flipped adenine bases and a cavity that could accommodate 2-DOS aminoglycoside binding (representative overlay with amikacin shown in Fig. 1G). These 12 homology-modeled *Bbu* ribosome structures were used to generate a total of 2088 (174×12) *Bbu* ribosome-MIA aminoglycoside complex structures.

**Figure 1.**
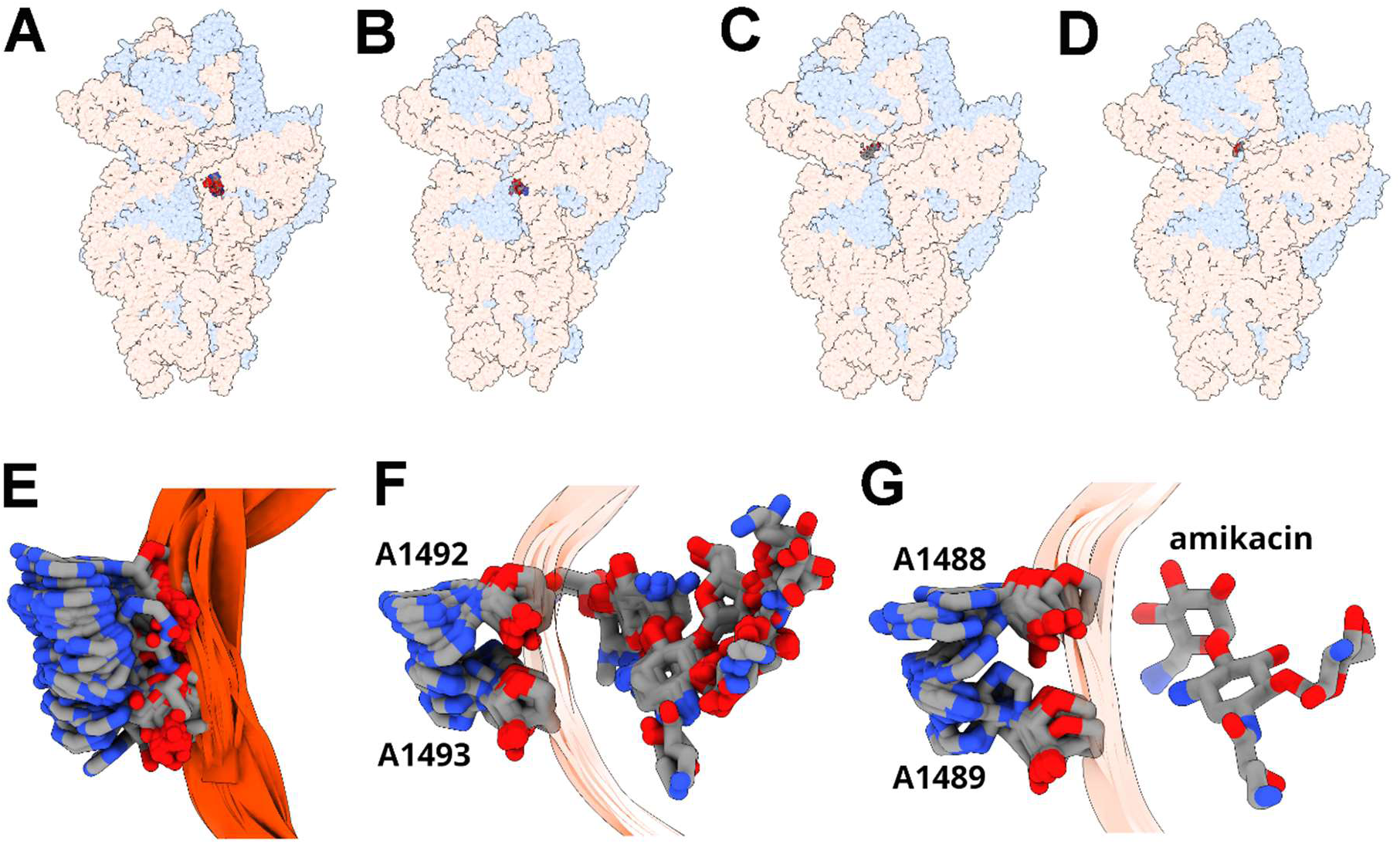
Four MIA binding sites in a bacterial 30S subunit; flipped conformation of conserved adenines that enable 2-DOS aminoglycoside binding; modeling of their representative conformations in *Bbu*. **(A)** 2-DOS aminoglycoside (paromomycin) overlay, **(A)** streptomycin overlay, **(A)** tetracycline overlay, **(A)** spectinomycin overlay, **(E)** overlay of flipped adenine conformations from all aminoglycoside-bound structures analyzed; **(F)** overlay of flipped adenine conformations for 12 highest resolution template structures bound to different aminoglycosides, **(G)** minimized structures of 12 homology models of the *Bbu* 30S subunit bound to amikacin. Ribosomal RNA is shown in orange red, ribosomal proteins in cornflower blue, and MIAs and the two conserved adenines are shown in stick format and are colored by element. The two conserved adenines are labeled by *Eco* numbering in panel **B** and by *Bbu* numbering in panel **C**.

**Table 5.** MIA non-aminoglycoside ribosome-bound structure details.

| Name | PDB ID | Res. (Å) | Method | Org. | RNA chain | Structure notes | Cit. |
| --- | --- | --- | --- | --- | --- | --- | --- |
| eravacycline (YQM) | 7M4U | 2.7 | Cryo-EM | *Aba* | a | 30S | 96 |
|  | 7M4V | 2.5 | Cryo-EM | *Aba* | A | 50S, **not used** | 96 |
|  | 7M4W | 2.6 | Cryo-EM | *Aba* | a | 70S | 96 |
|  | 7M4X | 2.7 | Cryo-EM | *Aba* | a | 70S | 96 |
|  | 7M4Y | 2.5 | Cryo-EM | *Aba* | a | 70S | 96 |
|  | 7M4Z | 2.9 | Cryo-EM | *Aba* | a | 70S, **not used** | 96 |
|  | 8CF8 | 2.2 | Cryo-EM | *Eco* | A | 30S | 45 |
| omadacycline (U3B) | 8CA7 | 2.1 | Cryo-EM | *Eco* | A | 30S | 45 |
| sarecycline (V7A) | 6XQD | 2.8 | X-ray | *Tth* | 1a, 2a | 70S | 97 |
|  | 6XQE | 3.0 | X-ray | *Tth* | 1a, 2a | 70S | 97 |
|  | 8CRX | 2.8 | Cryo-EM | *Cac* | A | 70S | 98 |
|  | 8CVM | 2.7 | Cryo-EM | *Cac* | a | 50S, **not used** | 98 |
|  | 8CVO | 3.0 | Cryo-EM | *Cac* | A | 30S | 98 |
| spectinomycin (SCM) | 1FJG | 3.0 | X-ray | *Tth* | A | 30S | 14 |
|  | 4V56 | 3.9 | X-ray | *Eco* | AA, CA | 70S | 48 |
|  | 4V57 | 3.5 | X-ray | *Eco* | AA, CA | 70S | 48 |
|  | 7N2V | 2.5 | Cryo-EM | *Eco* | 16 | 70S | 99 |
|  | 7P7Q | 2.4 | Cryo-EM | *Efa* | a | 70S | 100 |
|  | 7P7R | 2.9 | Cryo-EM | *Efa* | a | 70S | 100 |
|  | 7P7S | 3.0 | Cryo-EM | *Efa* | a | 70S | 100 |
|  | 7P7T | 2.9 | Cryo-EM | *Efa* | a | 70S | 100 |
|  | 7P7U | 3.1 | Cryo-EM | *Efa* | a | 70S | 100 |
|  | 7PIB | 4.7 | Cryo-EM | *Mpn* | 5 | 70S | 101 |
|  | 8CA7 | 2.1 | Cryo-EM | *Eco* | A | 30S | 45 |
|  | 8CAI | 2.1 | Cryo-EM | *Eco* | A | 30S | 45 |
|  | 8UVR | 2.6 | X-ray | *Tth* | 1a, 2a | 70S | 102 |
| tetracycline (TAC) | 1HNW | 3.4 | X-ray | *Tth* | A | 30S | 103 |
|  | 1I97 | 4.5 | X-ray | *Tth* | A | 30S | 104 |
|  | 4V9A | 3.3 | X-ray | *Tth* | AA, CA | 70S | 105 |
|  | 5J5B | 2.8 | X-ray | *Eco* | AA, BA | 70S | 106 |
|  | 5J7L | 3.0 | X-ray | *Eco* | AA, BA | 70S | 106 |
|  | 8CF1 | 1.8 | Cryo-EM | *Eco* | A | 30S | 45 |
|  | 8CGJ | 1.8 | Cryo-EM | *Eco* | A | 30S, **not used** | 45 |
| tigecycline (T1C) | 4V9B | 3.1 | X-ray | *Tth* | AA, CA | 70S | 105 |
|  | 4YHH | 3.4 | X-ray | *Tth* | A | 30S | 107 |
|  | 5J8A | 3.1 | X-ray | *Eco* | AA, BA | 70S | 106 |
|  | 5J91 | 3.0 | X-ray | *Eco* | AA, BA | 70S | 106 |
|  | 6YSI | 2.5 | Cryo-EM | *Aba* | 1 | 50S, **not used** | 42 |
|  | 6YTF | 3.0 | Cryo-EM | *Aba* | 3 | 30S | 42 |
|  | 8K2A | 2.9 | Cryo-EM | *Hsa* | S1 | 55S, **not used** | 108 |
|  | 8K2B | 3.4 | Cryo-EM | *Hsa* | L1 | 39S, **not used** | 108 |
|  | 8K2C | 2.4 | Cryo-EM | *Hsa* | S2 | 80S, **not used** | 108 |
|  | 8K2D | 3.2 | Cryo-EM | *Sce* | C2 | 80S, **not used** | 108 |
|  | 8K82 | 3.0 | Cryo-EM | *Sce* | C2 | 80S, **not used** | 108 |
|  | 8XSX | 2.4 | Cryo-EM | *Hsa* | S2 | 80S, **not used** | 108 |
|  | 8XSY | 3.0 | Cryo-EM | *Hsa* | S2 | 80S, **not used** | 108 |
|  | 8XSZ | 3.2 | Cryo-EM | *Hsa* | S2 | 80S, **not used** | 108 |
|  | 8XT0 | 3.2 | Cryo-EM | *Hsa* | S1 | 55S, **not used** | 108 |
|  | 8XT1 | 3.1 | Cryo-EM | *Hsa* | L1 | 39S, **not used** | 108 |
|  | 8XT2 | 3.3 | Cryo-EM | *Hsa* | S1 | 55S, **not used** | 108 |
|  | 8XT3 | 3.1 | Cryo-EM | *Hsa* | L1 | 39S, **not used** | 108 |
|  | 8YOO | 2.0 | Cryo-EM | *Hsa* | S2 | 80S, **not used** | 108 |

Res. = Resolution, Cit. = citation, np = no publication, RNA chain = chain(s) used to orient structures.

For MIAs that have no reported ribosomal small subunit-bound structure, the initial structure was either obtained from a non-ribosome-bound structure available from the RSCB Protein Databank or from predicted structures deposited in PubChem^31^. This initial structure was then fit as a rigid body into an artificial density map obtained using the molmap functionality of ChimeraX^29^ at a resolution of 1.5 Å applied to a high-resolution structure of a related MIA with a known structure that was already modeled in the *Bbu* ribosome. The 17 MIAs modeled in this fashion are astromicin, bekanamycin, dihydrostreptomycin, isepamicin, micronomicin, netilmicin, ribostamycin, chlortetracycline, clomocycline, demeclocycline, doxycycline, lymecycline, metacycline, minocycline, oxytetracycline, pipacycline, and rolitetracycline. The confidence in the prediction of proper binding conformations for these MIAs should be proportional to their chemical similarity to MIAs with known structures. For example, the confidence in *Bbu* ribosomal small subunit-bound structure for astromicin is lower because its binding site is not definitively known, while the predicted structure for dihydrostreptomycin has higher confidence given the amount of structural data available for modeling the binding of streptomycin.

### Single-point binding free energy estimations

All MIA aminoglycoside and non-aminoglycoside-bound *Bbu* ribosome structures generated were used to estimate MIA single-point binding free energies by Quickvina 2.1^32^. When we used re-docking of the MIA or unrestrained molecular dynamics simulations starting from an experimental structure, it was not possible to fully preserve the original pose of the MIA. To avoid altering the binding pose away from the experimentally determined starting position, neither docking nor molecular dynamics simulations were pursued. Instead, to consider variability in internal, positional, and environmental variables for binding of the MIAs to the *Bbu* ribosomal small subunit, we tested if starting MIA conformations could be restrictively minimized to improve their binding with minimal conformational change. For this minimization, the topology and parameters for each MIA were generated using CGENFF^33,34^ and the minimization was performed using CHARMM, version 40b2^35^. For the minimization, all atoms within 15 or 20 Å of the MIA were restrained using mass-weighted force constant of 1.0 kcal/(mol·Å²) and all atoms further away were held fixed. The minimization was performed using 5000 Steepest Descent (SD) steps and 500 Adopted-Basis Newton-Raphson steps to an energy tolerance of 0.001 kcal/mol. Single point binding free energies were estimated by Quickvina 2.1 for the original and the energy minimized structures. It should be noted that the energy functions used in CHARMM and Quickvina 2.1 are different, so the CHARMM minimization, which universally reduces the CHARMM-calculated energy, may not always improve the binding free energy estimated by Quickvina 2.1.

## Results and Discussion

### Aminoglycoside binding conformations

Amongst the MIAs binding to the ribosomal small subunit, the 2-DOS aminoglycosides have the most widespread clinical use. Of these, paromomycin has the largest number of deposited structures available (118, as listed in Table 3). Paromomycin likely helps stabilize tRNA binding and the orientation of the small subunit head region, which leads to better ribosome crystallization^36^. It is therefore included in ribosome structure determination protocols even when its own binding is not the purpose of the structure determination. The conformation of paromomycin and its environment in its highest resolution structure are shown in Fig. 2A and Fig. 2D, respectively. The large number of paromomycin structures can be used to obtain insights into the variability of its: (a) internal conformations (Fig. 2B), (b) positioning in the binding site (Fig. 2C), and (c) binding pocket residue conformations (Fig. 2E).

**Figure 2.**
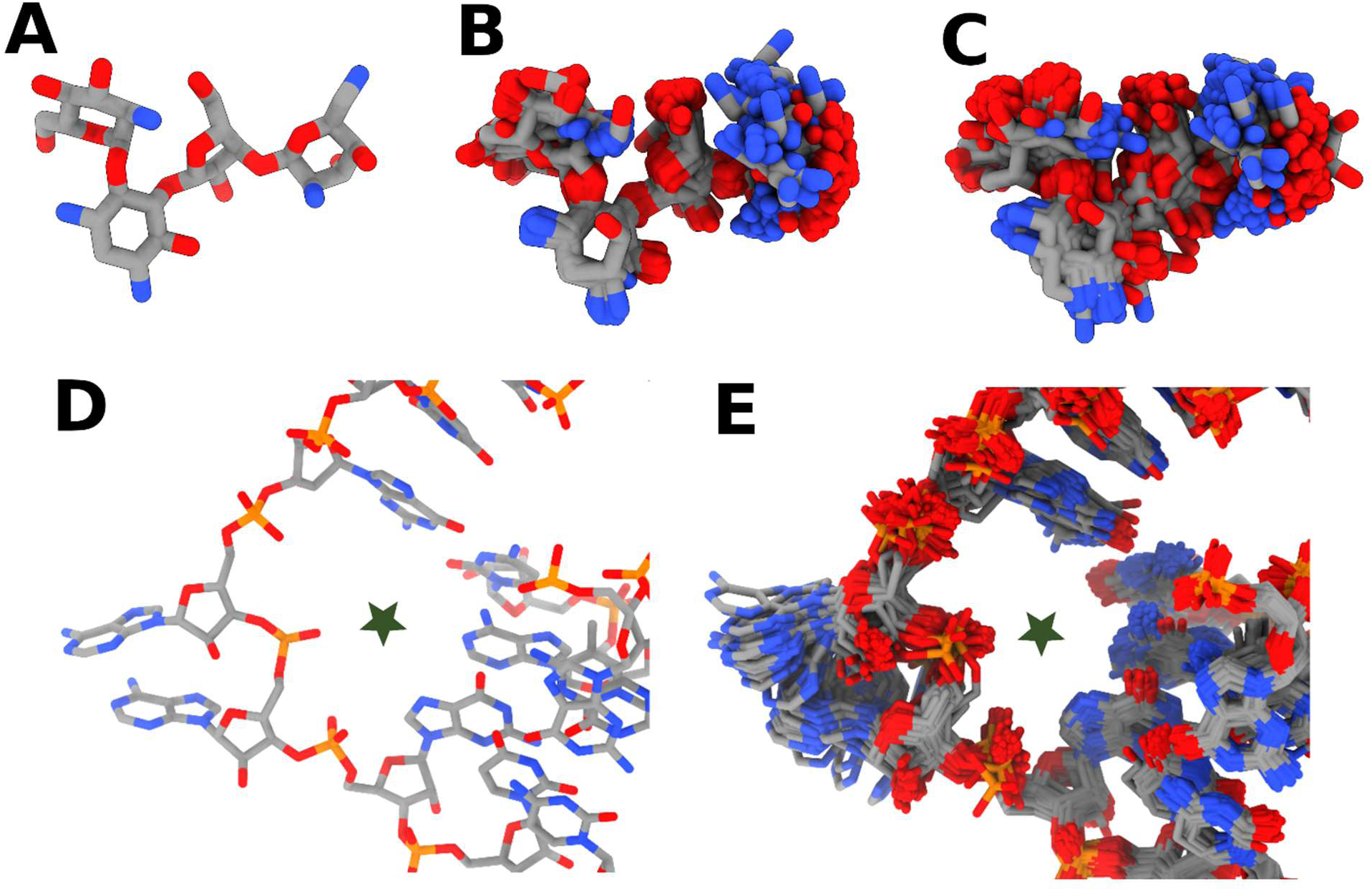
Conformational variability in paromomycin binding to ribosomal small subunits. **(A)** paromomycin conformation from the highest resolution structure (7K00^85^); **(B)** internal variability in paromomycin conformations, **(C)** positional variability for paromomycin in its binding pocket, **(D)** conformation of the surrounding 16S rRNA in the highest resolution structure (7K00^85^), **(E)** variability of the surrounding 16S rRNA in all paromomycin-bound structures. The location of paromomycin in panels D and E is indicated by a green star.

When self-aligned, the central 2-DOS and ribofuranosyl moieties of paromomycin appear less variable as compared to the outer idopyranosyl and glucopyranosyl moieties. When aligned according to the surrounding 16S rRNA, there seems to be additional paromomycin positional variability within the binding pocket (Fig. 2C). There is also variability in the conformation of the surrounding ribosomal small subunit binding region, especially the two flipped adenine bases (Fig. 2E). Since the resolution of the paromomycin-bound structures used in this analysis is between 2.0-4.5 Å, it is likely that the observed variability in the paromomycin and surrounding residue conformations is not due to modeling errors. A movie showing transitions between these different paromomycin structures illustrates this variability visually (movie S1). There is also internal and positional variability detectable in other aminoglycosides that have more than one structure, when they are bound to the ribosomal small subunit. This can be seen in the overlays generated by aligning their structures using the surrounding 16S rRNA (Fig. 3A-L). This variability is not just for the 2-DOS aminoglycoside binding pocket formed by flipping of the two adenine bases, it is also seen for streptomycin (Fig. 3M, movie S2), which is chemically distinct and binds to a different 30S pocket.

**Figure 3.**
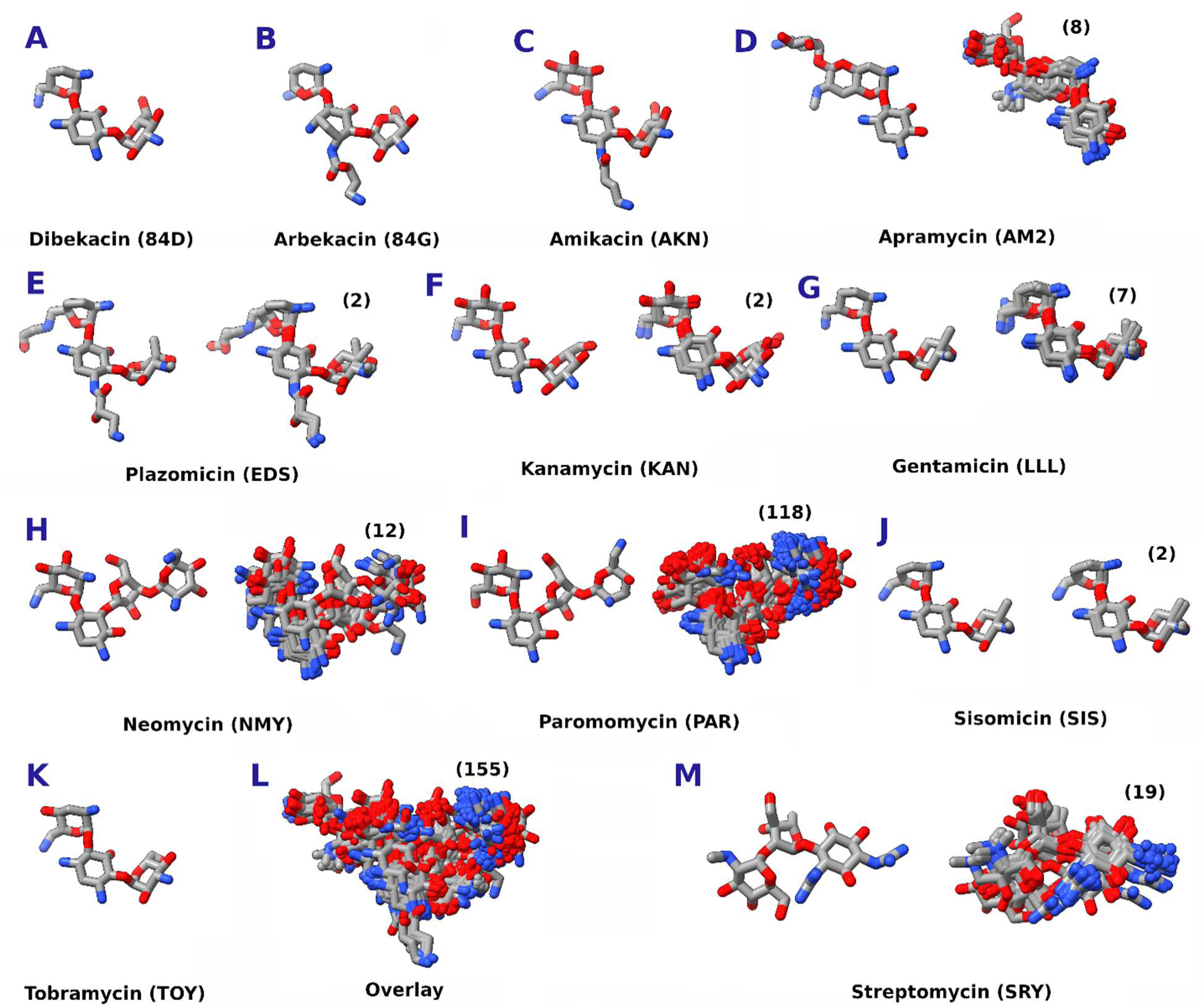
Conformational variability of aminoglycoside MIAs bound to ribosomal small subunits. **(A)** dibekacin, **(B)** arbekacin, **(C)** amikacin, **(D)** apramycin, **(E)** plazomicin, **(F)** kanamycin, **(G)** gentamicin, **(H)** neomycin, **(I)** paromomycin, **(J)** sisomicin, **(K)** tobramycin, **(L)** Overlay of all 2-DOS aminoglycosides, **(M)** streptomycin. For aminoglycosides with more than one structure, the highest resolution structure is shown on the left and an overlay obtained by aligning their surrounding 16S rRNA is shown on the right. The number of structures for each individual aminoglycoside is shown in parentheses.

### Non-aminoglycoside binding conformations

Tetracycline analogs are a clinically important class of non-aminoglycoside MIAs that bind to the ribosomal small subunit^37^. Their internal conformations and positional variability in their 30S binding pocket are shown in Fig. 4A-F. The number of structures of tetracycline and its analogs bound to their expected functional bacterial ribosomal small subunit binding site are fewer (a total of 29 structures, Fig. 4F) than those for aminoglycosides, but they also show internal and positional variability for their binding conformations. This observation is also consistent for spectinomycin, another non-aminoglycoside MIA that has 16 individual ribosomal small subunit-bound structures (Fig. 4G). For both aminoglycosides and non-aminoglycosides binding to the ribosomal small subunit, the internal and positional variability seems to increase as the number of available structures for a particular MIA increase. This suggests that a single structure may not capture the possible variability for that MIA or its binding pocket.

**Figure 4.**
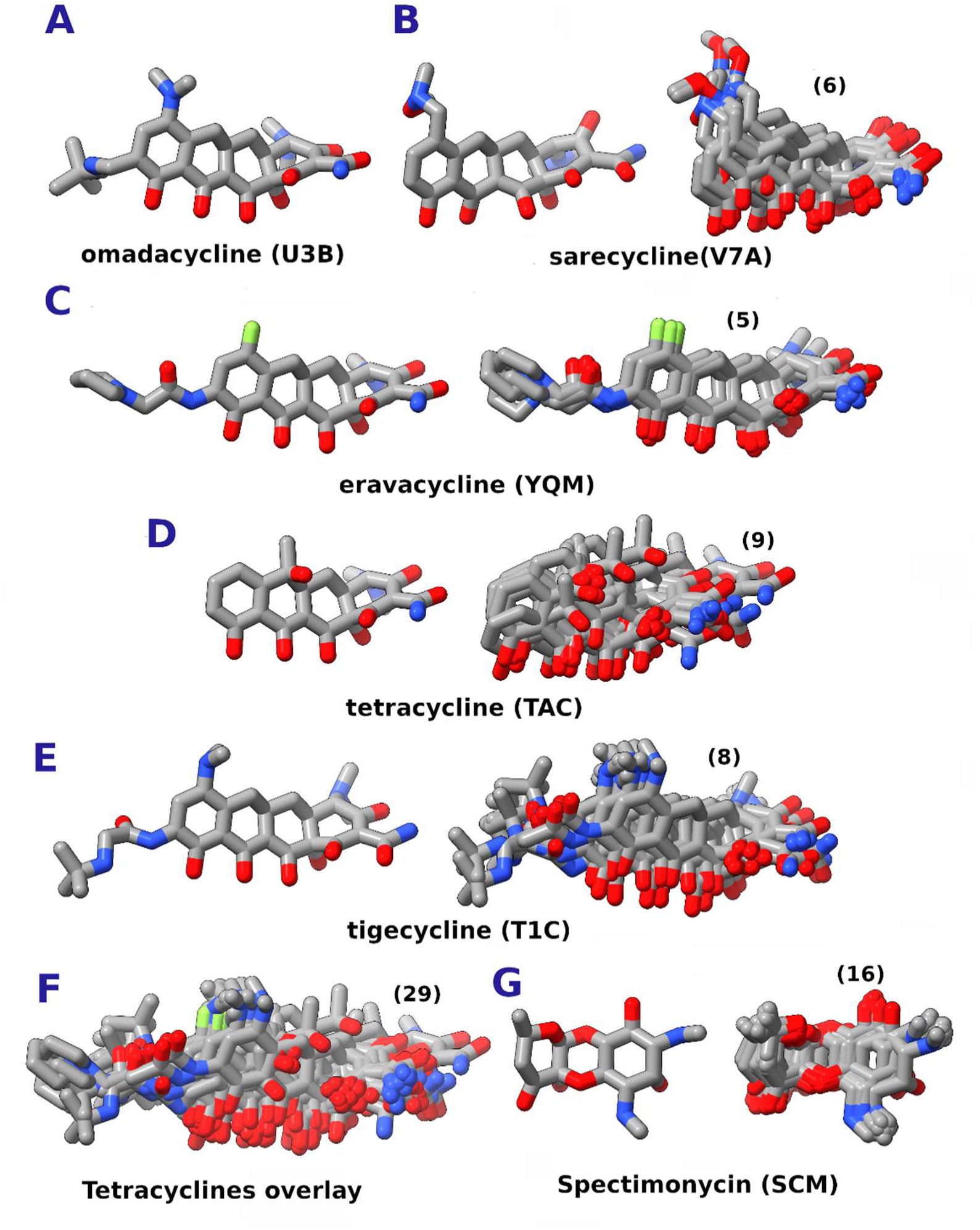
Conformational variability of non-aminoglycoside MIAs bound to ribosomal small subunits. **(A)** omadacycline, **(B)** sarecycline, **(C)** eravacycline, **(D)** tetracycline, **(E)** tigecycline, **(F)** tetracycline analog overlay, **(G)** spectinomycin. For non-aminoglycosides with more than one structure, the highest resolution structure is shown on the left and an overlay obtained by aligning their surrounding 16S rRNA is shown on the right. The number of structures for each individual non-aminoglycoside is shown in parentheses.

Individual experimentally determined structures have composition differences, e.g. in the sequences of the ribosomal small subunit components or in the presence of tRNAs or external protein factors, but these were not necessarily thought to significantly affect MIA binding. The internal and positional variability of the MIA structures shows that these factors may be important in the precise conformation adopted by the MIA and therefore also its binding affinity in those conformations. Evolutionary analysis shows that sites that bind ribosomal inhibitors can be divergent across species^38^. Structure-based design of new antibiotics therefore should make choices for synthetic MIA modifications by accounting for both the structural variability of the MIA and its environment.

### Prediction of MIA binding to the *Bbu* 30S subunit

Structural models for the *Bbu* ribosome bound to all MIAs listed in Table 1 were generated as described in the methods. To account for variability in flipped conformations of conserved adenines A1488 and A1489 (*Bbu* numbering) during aminoglycoside MIA binding, 12 representative conformations of residues 1485-1492 were modeled. These modeled structures were then gently minimized to maintain the structure of the environment and the MIA as close as possible to their original structures. The resulting models were then analyzed for their single-point binding free energy using Quickvina 2.1^32^ and these binding free energies were sorted to find the conformation with the best binding free energy, which is to be considered the optimal binding conformation predicted using this approach. For the MIAs with previously known structures bound to bacterial ribosomes, the best binding free energy conformations are shown in Fig. 5 for the aminoglycoside MIAs and in Fig. 6 for the non-aminoglycoside MIAs. The overlays of all modeled bound conformations obtained from previously known structures are shown in Fig. S1 for the aminoglycoside MIAs and in Fig. S2 for the non-aminoglycoside MIAs. The overlays of minimized conformations for MIAs with no previous structures bound to a 30S subunit are shown in Fig. S3. For these MIAs, the confidence in the structural predictions (shown in Fig. 7) should be proportional to their similarity to previously known MIAs. Experimental structural determination for these MIAs with no known structures would confirm their binding conformations to the bacterial ribosome and help validate or improve our computational structure prediction approach for antibiotic binding.

**Figure 5.**
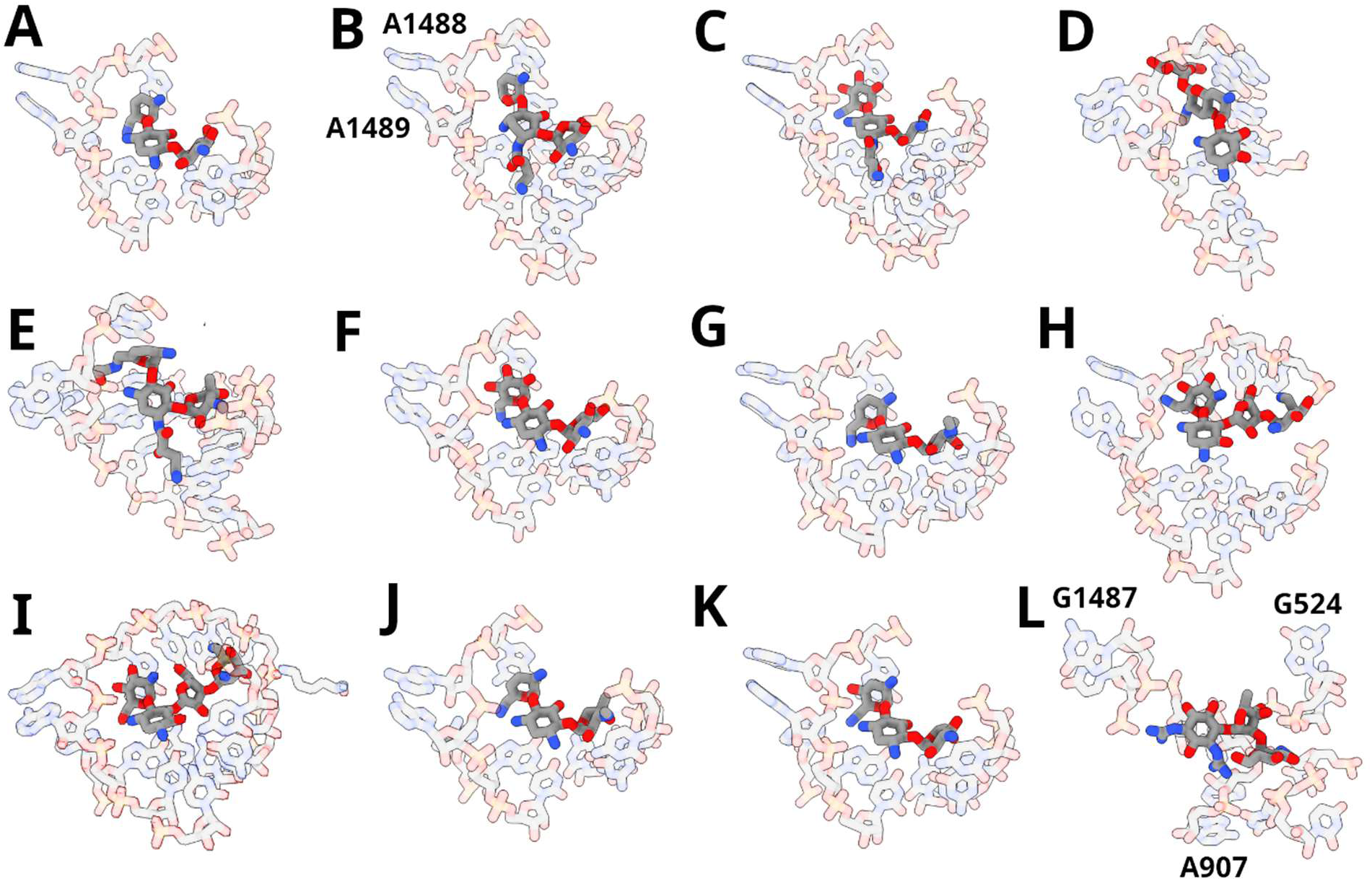
Best binding free energy predicted conformations of *Bbu* 30S-bound complexes for aminoglycoside MIAs with existing ribosome-bound experimental structures. **(A)** dibekacin, **(B)** arbekacin, **(C)** amikacin, **(D)** apramycin, **(E)** plazomicin, **(F)** kanamycin, **(G)** gentamicin, **(H)** neomycin, **(I)** paromomycin, **(J)** sisomicin, **(K)** tobramycin, **(L)** streptomycin. MIA structures are shown in opaque stick format while the residues within 3 Å of any MIA atom in its environment are shown in transparent stick format.

**Figure 6.**
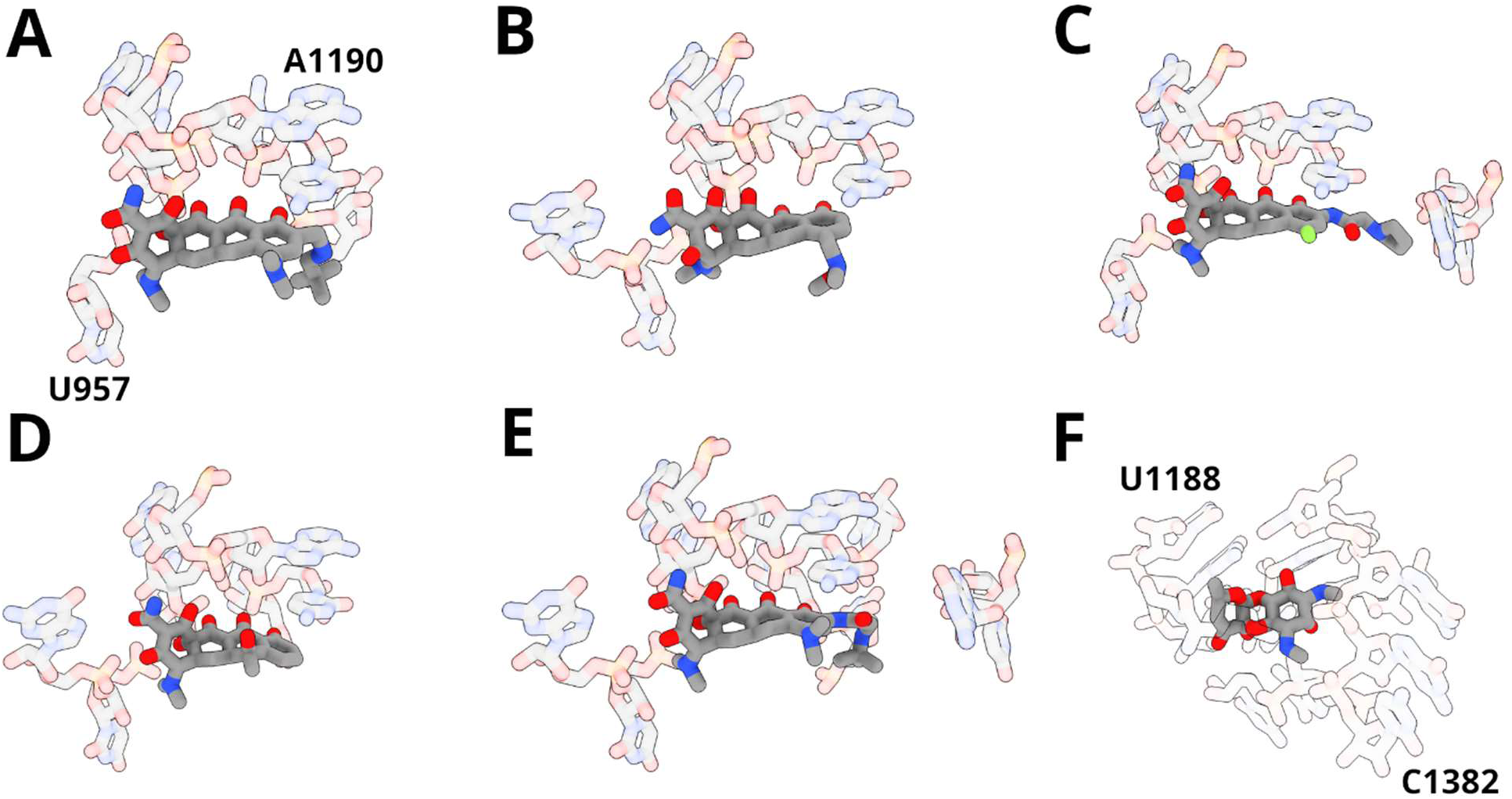
Best binding free energy structural predictions of *Bbu* 30S-bound complexes for non-aminoglycoside MIAs with existing ribosome-bound experimental structures. **(A)** omadacycline, **(B)** sarecycline, **(C)** eravacycline, **(D)** tetracycline, **(E)** tigecycline, **(F)** spectinomycin. MIAs are shown in opaque stick format while the residues within 3 Å of any MIA atom in its environment are shown in transparent stick format.

**Figure 7.**
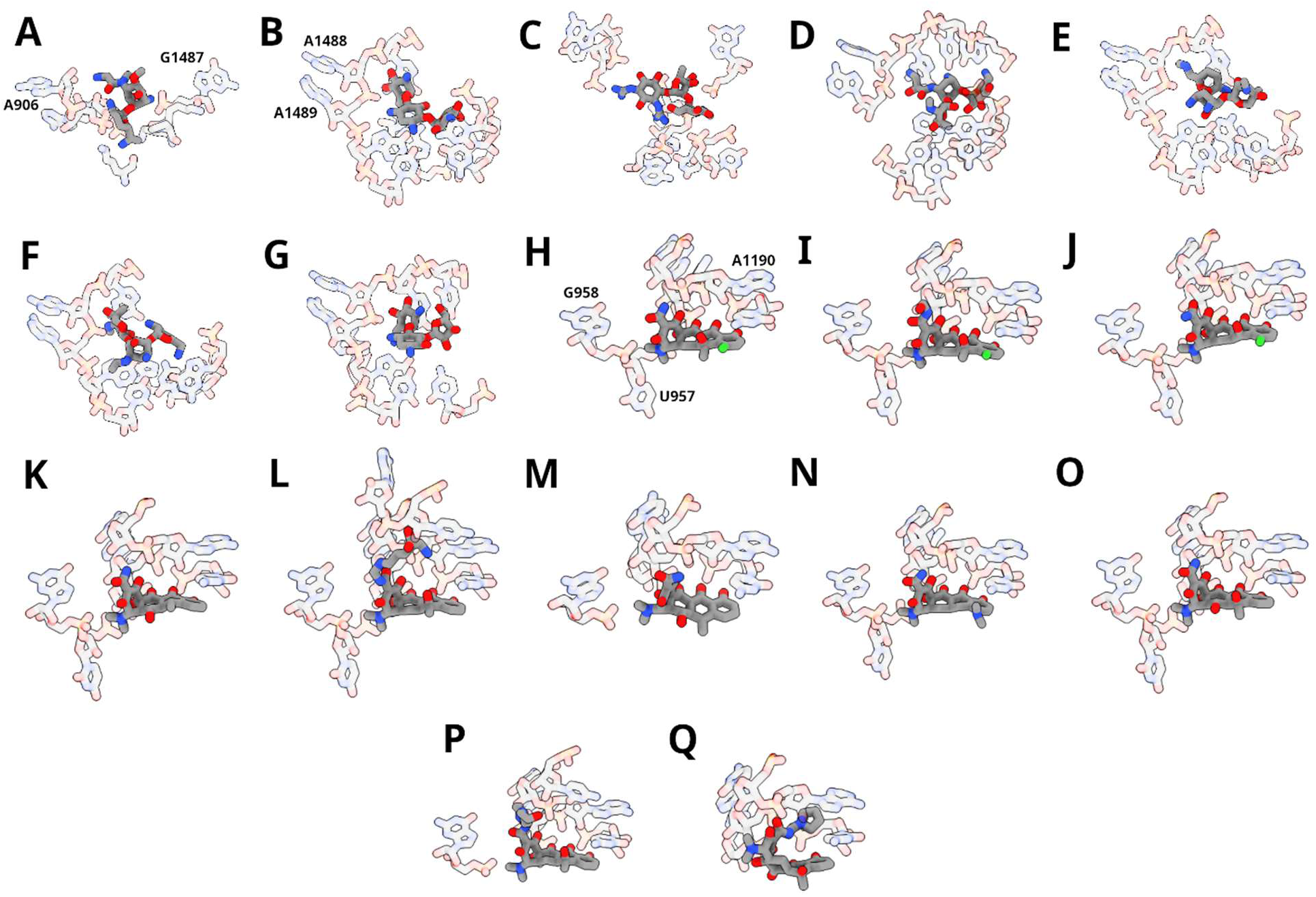
Best binding free energy structural predictions of *Bbu* 30S-bound complexes for MIAs with no previous ribosome-bound experimental structure. **(A)** astromicin, **(B)** bekanamycin, **(C)** dihydrostreptomycin, **(D)** isepamicin, **(E)** micronomicin, **(F)** netilmicin, **(G)** ribostamycin, **(H)** chlortetracycline, **(I)** clomocycline, **(J)** demeclocycline, **(K)** doxycycline, **(L)** lymecycline, **(M)** metacycline, **(N)** minocycline, **(O)** oxytetracycline, **(P)** pipacycline, and **(Q)** rolitetracycline. MIAs are shown in opaque stick format while the residues within 3 Å of any MIA atom in its environment are shown in transparent stick format.

### Free energies for MIA binding to the *Bbu* 30S subunit

The single-point binding free energies for minimized models of *Bbu* 30S subunit-bound MIAs with previously known structures are listed in Table 6. The difference between the minima and maxima, and the standard deviations of these single-point binding free energies point to their sensitivity to the specific initial conformation of the MIA. For example, for the 144 conformations modeled for neomycin, the minimum value is -6.1 kcal/mol whereas the maximum value is +2.9 kcal/mol. In the worst case within the parameters of the present approach, a binding free energy prediction made from a minimized model of a single experimentally derived structural pose of neomycin could therefore have an error up to 9 kcal/mol. Since each of these minimized models is obtained from an experimentally determined MIA structure, predicting an optimal model from a single known MIA structure clearly involves some risk.

**Table 6.** *Bbu* 30s subunit binding free energies for MIAs with known structures.

| **MIA** | **Minimum BFE** | **Average BFE** | **Maximum BFE** | **BFE Std. Dev.** | **Number of models** |
| --- | --- | --- | --- | --- | --- |
| apramycin | -6.8 | -5.2 | -0.1 | 1.0 | 96 |
| paromomycin | -6.5 | -4.7 | -2.1 | 0.8 | 1416 |
| dibekacin | -6.3 | -5.8 | -5.0 | 0.4 | 12 |
| neomycin | -6.1 | -2.6 | 2.9 | 3.0 | 144 |
| tobramycin | -6.1 | -5.8 | -5.4 | 0.2 | 12 |
| gentamicin | -5.9 | -5.1 | -3.5 | 0.5 | 84 |
| kanamycin | -5.6 | -5.0 | -4.3 | 0.4 | 24 |
| amikacin | -5.4 | -4.6 | -4.1 | 0.4 | 12 |
| sisomicin | -5.3 | -4.1 | -2.4 | 0.8 | 24 |
| spectinomycin | -5.2 | -4.5 | -3.3 | 0.4 | 16 |
| tetracycline | -5.0 | -2.4 | 0.8 | 2.0 | 9 |
| tigecycline | -4.4 | -2.0 | 0.3 | 2.0 | 8 |
| omadacycline | -4.4 | -4.4 | -4.4 | 0.0 | 1 |
| sarecycline | -4.2 | -2.4 | -1.2 | 1.0 | 6 |
| plazomicin | -4.2 | -2.9 | -1.2 | 0.9 | 24 |
| arbakacin | -4.1 | -3.6 | -2.9 | 0.4 | 12 |
| streptomycin | -3.8 | -1.9 | 0.0 | 0.9 | 246 |
| eravacycline | -3.6 | -2.4 | -1.6 | 0.7 | 5 |

BFE: Single point binding free energy calculated by Quickvina2.1 (in kcal/mol), Std. Dev.: Standard deviation

In the present study, a gentle minimization is carried out using the CHARMM energy function^35^ while the binding free energy is estimated using the Autodock Vina energy function^39^. Comparison of single-point binding free energies computed before and after minimization for paromomycin (Fig. S4A) and streptomycin (Fig. S5A) show that even this very restrained minimization can improve the binding free energies substantially. The structures for the conformations before and after minimization can also be very similar even when the change in the binding free energy is large, i.e. the conformational changes required to significantly improve the binding free energy can be small, as seen in the examples for paromomycin (Fig. S4B-D) and streptomycin (Fig. S5B-D).

The binding free energies for the optimal MIA models bound to the *Bbu* 30S subunit range from -6.8 kcal/mol (apramycin) to -3.6 kcal/mol (eravacycline). These single-point binding free energies are calculated with a crude empirical model^39^ consisting of pair-wise atomic terms for attractive, repulsive, hydrophobic, and hydrogen-bonding interactions, and a torsional penalty, and are therefore prone to internal errors. It would not be prudent, therefore, to suggest apramycin as a clinical MIA of choice for Lyme disease simply based on its predicted favorable binding free energy. More sophisticated and detailed energy functions could reduce such errors but are expected to have a similar sensitivity to the specific MIA starting conformation. One way to resolve the problem computationally would be to calculate absolute binding free energies^40^ using explicit solvent molecular dynamics simulations with very detailed energy functions^41^ but these will also be susceptible to other issues such as high computational cost, insufficient sampling, and possible inaccuracies in the composition of post-transcriptional or post-translational modifications, ions, or solvent in the environment. If all such issues can be resolved satisfactorily, however, it should be possible to anticipate the best MIA for a particular bacterial ribosome, which could enable species-specific narrow-spectrum therapeutics that could minimize the effect on the rest of the bacterial microbiome.

## Conclusions

Understanding the detailed structural and physical basis of MIA binding to the ribosomal small subunit will enable fast and inexpensive design of the next generation of 30S-targeting antibiotics based on existing MIA scaffolds. The present work builds upon previous structural data of MIA binding to bacterial 30S subunits to predict MIA binding to the *Bbu* 30S subunit. This provides the first compilation of optimized models for all MIAs bound to the bacterial 30S subunit of a single bacterial species (movie S3). We are now applying the present protocol to generate optimized models of all MIAs bound to the *Bbu* ribosomal large 50S subunit. Our inexpensive computational approach can then be extended to anticipate MIA binding to full ribosomes of all other bacterial pathogen species.

The starting conformation sensitivity of the computed binding free energies for MIA binding to the *Bbu* 30S subunit suggests that accuracy in the structural details of MIA binding is critical. Small changes in the MIA structure or the structure and composition of its environment can have a large impact on its binding free energy. MIAs with no known experimentally determined 30S-bound structure therefore represent a significant knowledge gap that should be filled through experimental efforts. It is also likely that the binding free energy of individual MIAs to different bacterial ribosomes is not identical. Here, we have assessed MIA binding to an important pathogen ribosome, but a wider assessment of binding free energies and experimentally determined binding affinities to different pathogen ribosomes is a necessary subsequent step.

This approach is directly applicable to anticipate changes in MIA binding due to resistance mutations in bacterial ribosomes through calculation of MIA binding free energies in the presence and absence of the resistance mutations. It can be used to understand the selectivity of specific MIAs for eukaryotic pathogen ribosomes by predicting the binding free energies for MIAs in those ribosomes. MIA binding to host cytosolic and organellar ribosomes can also be predicted, providing a way to understand the basis of MIA toxicity. This can also provide a pathway to ameliorate MIA toxicity through structurally guided chemical modifications that reduce binding to host ribosomes while maintaining or improving binding to pathogen ribosomes.

## Supporting information

CHARMM topology/parameters for MIAs

Optimized MIA models bound to ribosome

## Author contributions

NKB designed the project; SRM, CM, and NKB performed the research, SRM, CM, and NKB wrote and edited the manuscript.

## Acknowledgments

Feedback from Dr. Rajendra Agrawal and his group members was helpful to this work.

## Supplementary material

Five figures (Fig. S1-S5), three movies (movies S1-S3), a zip file with CHARMM format topology and parameter files for all MIAs modeled, and a pdb file with optimized models of 35 MIAs with their neighboring environment in the *Bbu* ribosome and a minimal protein backbone and nucleic acid phosphorus atom model of the *Bbu* 30S subunit from the PDB deposition (PDB ID: 8FMW) are included as supplementary material.

## Movie captions

**Movie S1.** Transitions between different structures of paromomycin bound to ribosomes. Paromomycin is shown in green while ribosomal residues within 5 Å of paromomycin are shown in element colors in transparent stick format.

**Movie S2.** Transitions between different structures of streptomycin bound to ribosomes. Streptomycin is shown in green while ribosomal residues within 5 Å of streptomycin are shown in element colors in transparent stick format.

**Movie S3.** Most favorable binding free energy minimized models of 35 MIAs binding to the *Bbu* 30S subunit. MIAs shown in element color and stick format, RNA shown in orange red and proteins shown in cornflower blue.

