## Supplementary material pdf for "How medically important antimicrobials bind to the 30S ribosomal subunit in a bacterial pathogen"

Five figures (Fig. S1-S5), three movies (movies S1-S3), a zip file with CHARMM format topology and parameter files for all MIAs modeled, and a pdb file with optimized models of 35 MIAs with their neighboring environment in the *Bbu* ribosome and a minimal protein backbone and nucleic acid phosphorus atom model of the *Bbu* 30S subunit from the PDB deposition (PDB ID: 8FMW) are included as supplementary material.

**Movie captions.**

**Movie S1.** Transitions between different structures of paromomycin bound to ribosomes. Paromomycin is shown in green while ribosomal residues within 5 Å of paromomycin are shown in element colors in transparent stick format.

**Movie S2.** Transitions between different structures of streptomycin bound to ribosomes. Streptomycin is shown in green while ribosomal residues within 5 Å of streptomycin are shown in element colors in transparent stick format.

**Movie S3.** Most favorable binding free energy minimized models of 35 MIAs binding to the *Bbu* 30S subunit. MIAs shown in element color and stick format, RNA shown in orange red and proteins shown in cornflower blue.

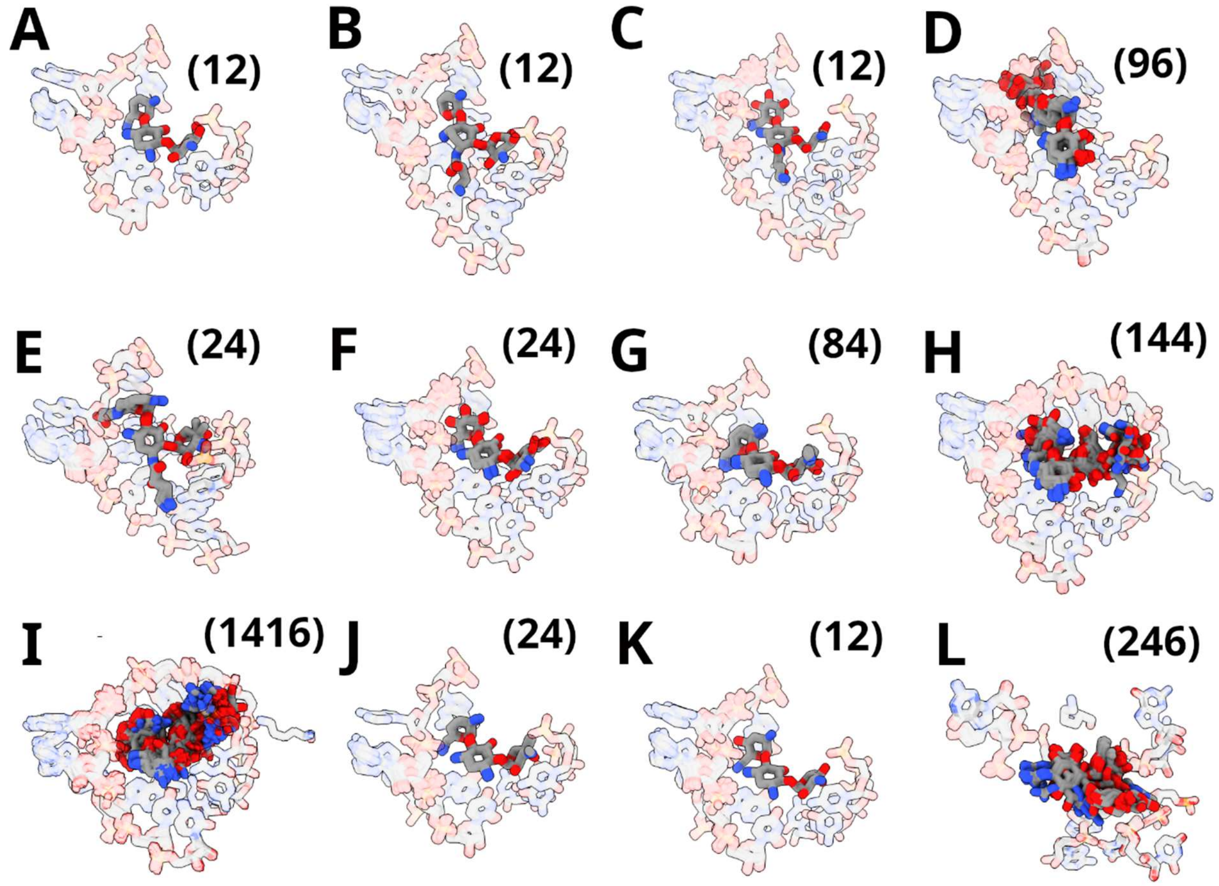


**Figure S1. Overlay of structural predictions of *Bbu* 30S-bound complexes for aminoglycoside MIAs with existing ribosome-bound experimental structures. (A)** dibekacin, **(B)** arbekacin, **(C)** amikacin, **(D)** apramycin, **(E)** plazomicin, **(F)** kanamycin, **(G)** gentamicin, **(H)** neomycin, **(I)** paromomycin, **(J)** sisomicin, **(K)** tobramycin, **(L)** streptomycin. MIA structures are shown in opaque stick format while the residues within 3 Å of any MIA atom in its environment are shown in transparent stick format. Numbers in parentheses represent number of overlaid *Bbu* 30S-bound MIA models shown.

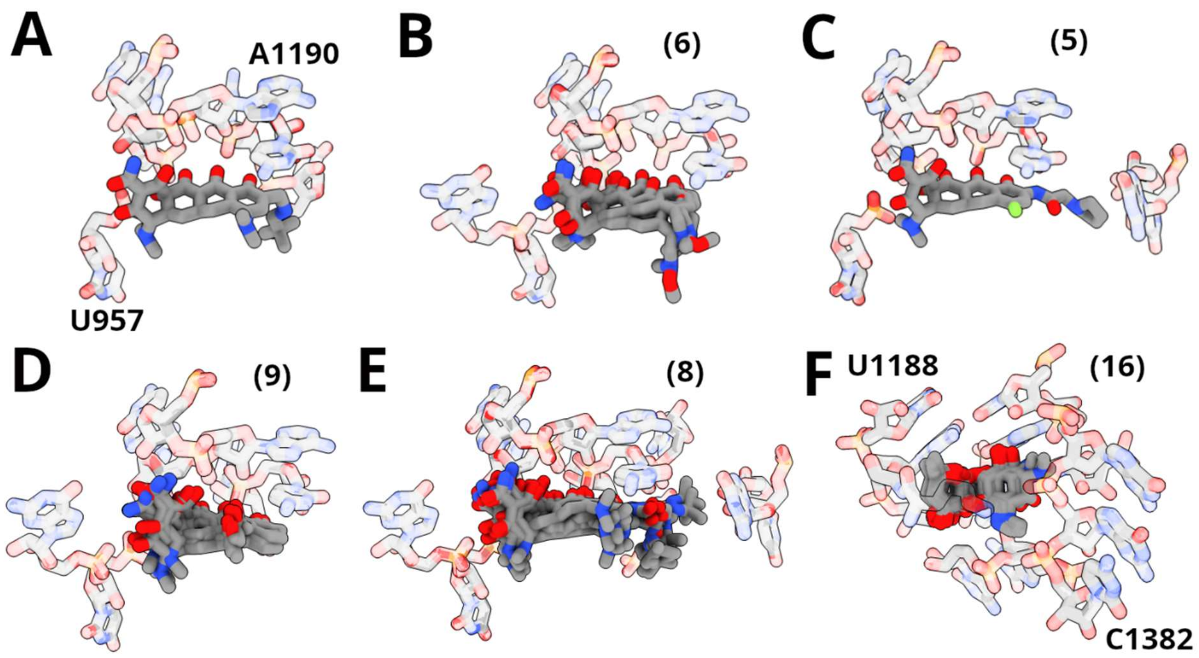


**Figure S2. Overlay of structural predictions of *Bbu* 30S-bound complexes for non-aminoglycoside MIAs with existing ribosome-bound experimental structures. (A)** omadacycline, **(B)** sarecycline, **(C)** eravacycline, **(D)** tetracycline, **(E)** tigecycline, **(F)** spectinomycin. MIAs are shown in opaque stick format while the residues within 3 Å of any MIA atom in its environment are shown in transparent stick format. Numbers in parentheses represent number of overlaid *Bbu* 30S-bound MIA models shown.

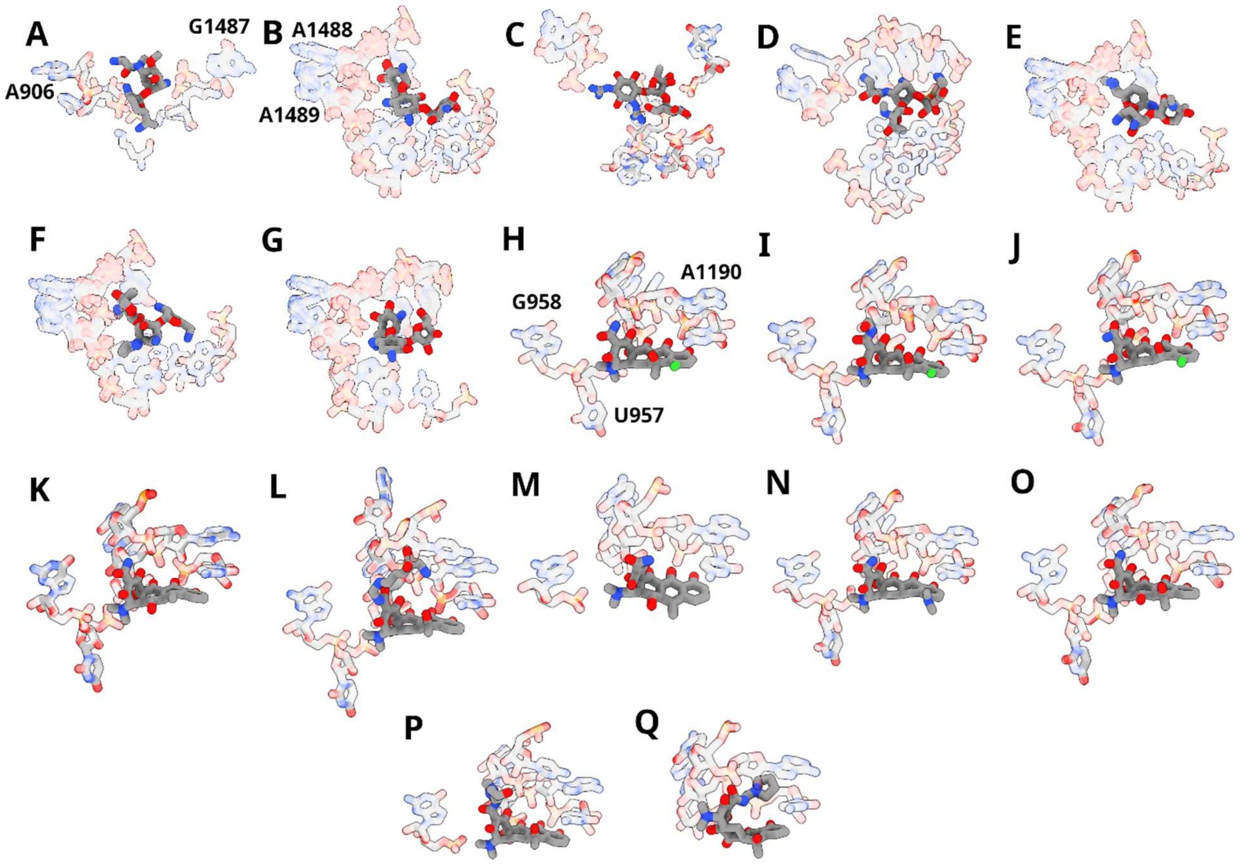


**Figure S3. Overlay of structural predictions of *Bbu* 30S-bound complexes for MIAs with no previous ribosome-bound experimental structure. (A)** astromicin, **(B)** bekanamycin, **(C)** dihydrostreptomycin, **(D)** isepamicin, **(E)** micronomicin, **(F)** netilmicin, **(G)** ribostamycin, **(H)** chlortetracycline, **(I)** clomocycline, **(J)** demeclocycline, **(K)** doxycycline, **(L)** lymecycline, **(M)** metacycline, **(N)** minocycline, **(O)** oxytetracycline, **(P)** pipacycline, and **(Q)** rolitetracycline. MIAs are shown in opaque stick format while the residues within 3 Å of any MIA atom in its environment are shown in transparent stick format.

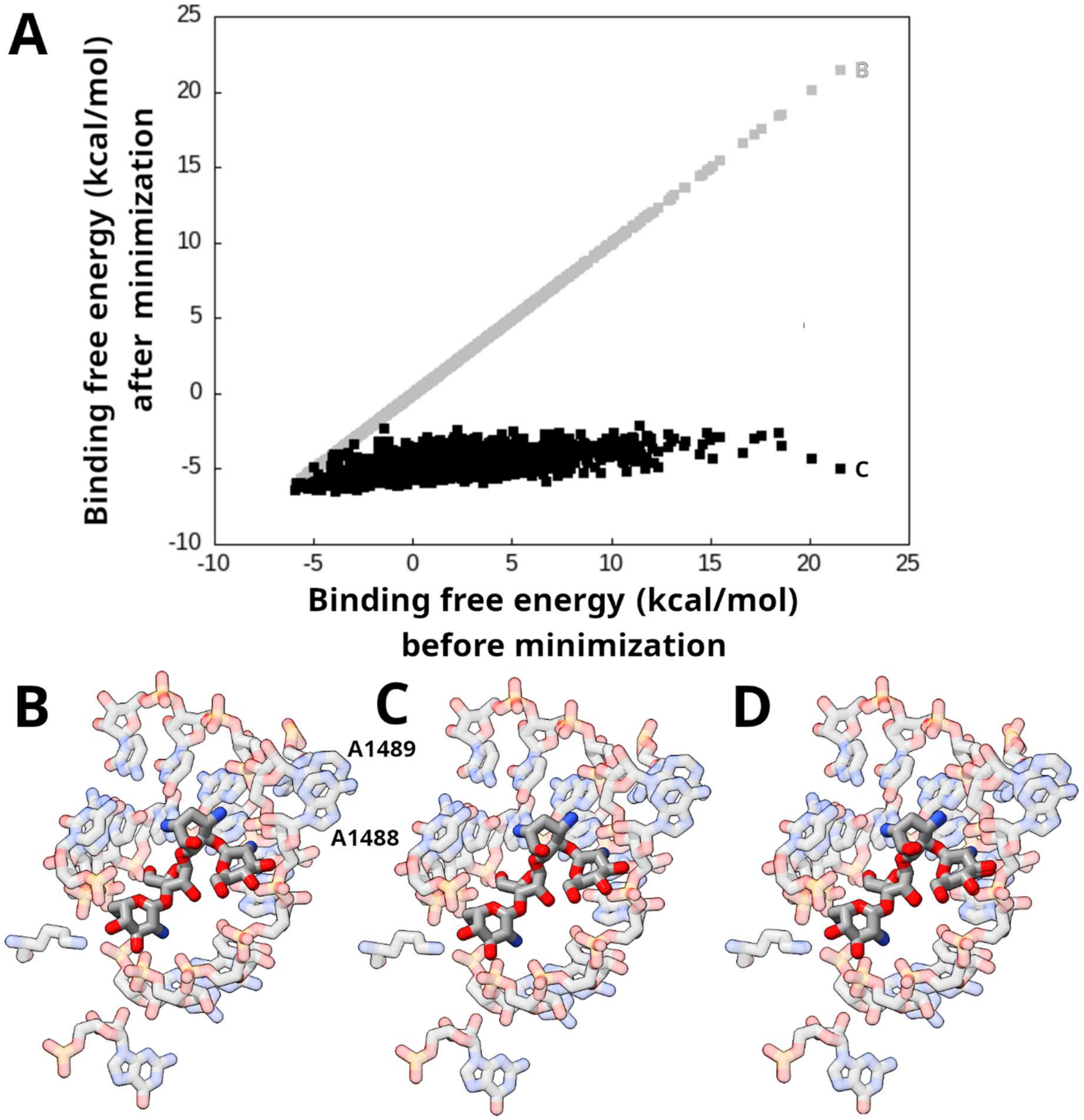


**Figure S4. Effect of minimization on single point binding free energies for paromomycin. (A)** Scatter plot of binding free energies before and after minimization for all paromomycin - *Bbu* 30S subunit complexes modeled. Grey diagonal values depict the scatter plot if there was no change in binding free energy due to minimization. **(B)** An unfavorable binding free energy conformation before minimization, **(C)** The favorable binding free energy structure after minimization of conformation in panel B, **(D)** Overlay of the conformations in panels **B** and **C**. Paromomycin shown in opaque sticks while residues within 5 Å of paromomycin shown in transparent sticks. Binding free energies of conformations in panels **B** and **C** are labeled in panel **A**.

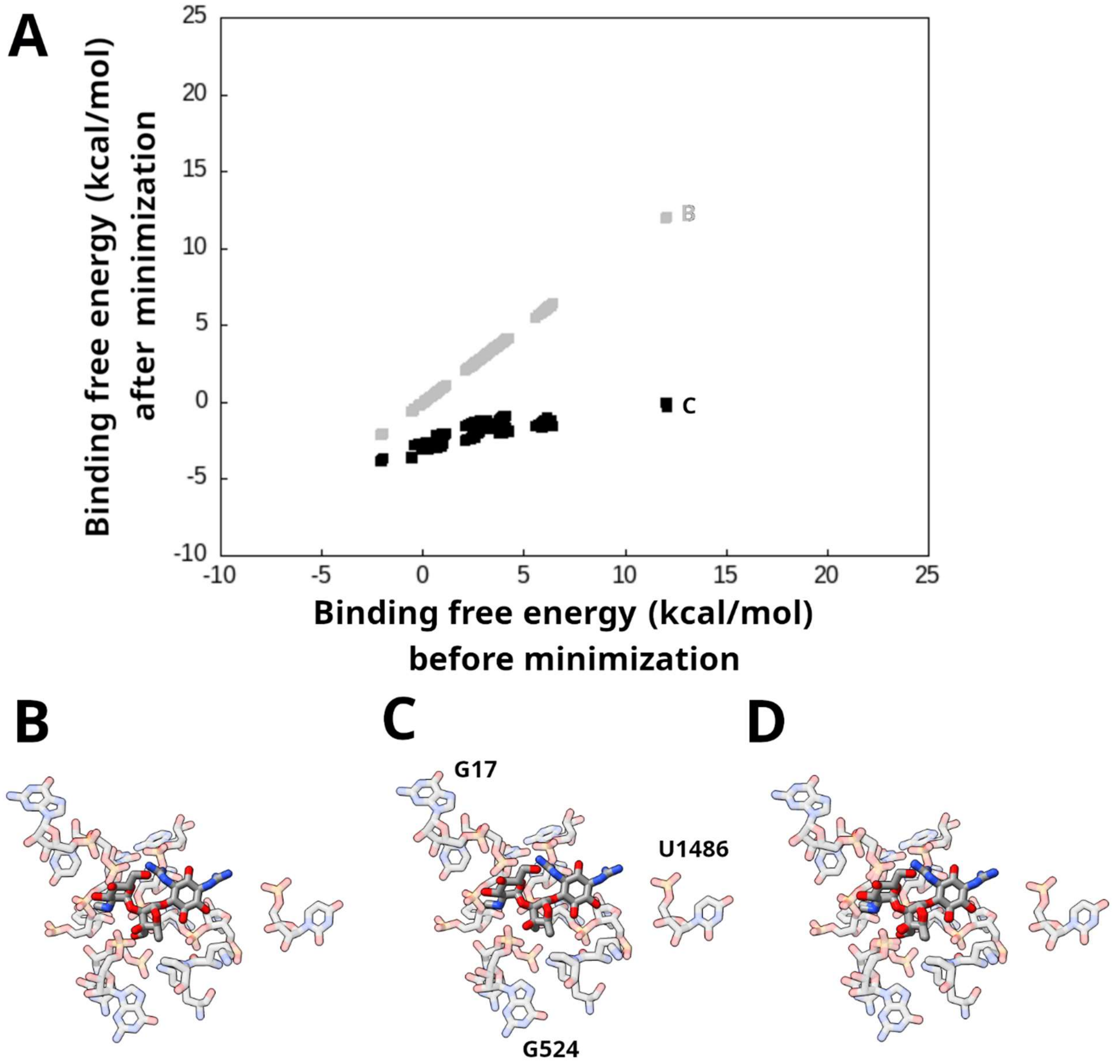


**Figure S5. Effect of minimization on single point binding free energies for streptomycin. (A)** Scatter plot of binding free energies before and after minimization for all streptomycin - *Bbu* 30S subunit complexes modeled. Grey diagonal values depict the scatter plot if there was no change in binding free energy due to minimization. **(B)** An unfavorable binding free energy conformation before minimization, **(C)** The favorable binding free energy structure after minimization of conformation in panel B, **(D)** Overlay of the conformations in panels **B** and **C**. Streptomycin shown in opaque sticks while residues within 5 Å of streptomycin shown in transparent sticks. Binding free energies of conformations in panels **B** and **C** are labeled in panel **A**.
