## Supplementary material for "How medically important antimicrobials bind to the 30S ribosomal subunit in a bacterial pathogen": CHARMM topology/parameters for MIAs: 35mia_30s_cgenff_toppar_readme.txt

Supplementary Material for:
'How medically important antimicrobials bind to the 30S ribosomal subunit in a bacterial pathogen'
Swati R. Manjari, Caleb Mallery, Nilesh K. Banavali

This gzipped tar file contains the following files for 35 medically important antimicrobials (MIAs) that
can be used for modeling them in CHARMM.

amikacin_cgenff_toppar.str: amikacin CHARMM format topology and parameters, requires cgenff parameters to be read in before
apramycin_cgenff_toppar.str: apramycin CHARMM format topology and parameters, requires cgenff parameters to be read in before
arbekacin_cgenff_toppar.str: arbekacin CHARMM format topology and parameters, requires cgenff parameters to be read in before
astromicin_cgenff_toppar.str: astromicin CHARMM format topology and parameters, requires cgenff parameters to be read in before
bekanamycin_cgenff_toppar.str: bekanamycin CHARMM format topology and parameters, requires cgenff parameters to be read in before
chlortetracycline_cgenff_toppar.str: chlortetracycline CHARMM format topology and parameters, requires cgenff parameters to be read in before
clomocycline_cgenff_toppar.str: clomocycline CHARMM format topology and parameters, requires cgenff parameters to be read in before
demeclocycline_cgenff_toppar.str: demeclocycline CHARMM format topology and parameters, requires cgenff parameters to be read in before
dibekacin_cgenff_toppar.str: dibekacin CHARMM format topology and parameters, requires cgenff parameters to be read in before
dihydrostreptomycin_cgenff_toppar.str: dihydrostreptomycin CHARMM format topology and parameters, requires cgenff parameters to be read in before
doxycycline_cgenff_toppar.str: doxycycline CHARMM format topology and parameters, requires cgenff parameters to be read in before
eravacycline_cgenff_toppar.str: eravacycline CHARMM format topology and parameters, requires cgenff parameters to be read in before
framycetin_cgenff_toppar.str: framycetin CHARMM format topology and parameters, requires cgenff parameters to be read in before
gentamicin_cgenff_toppar.str: gentamicin CHARMM format topology and parameters, requires cgenff parameters to be read in before
isepamicin_cgenff_toppar.str: isepamicin CHARMM format topology and parameters, requires cgenff parameters to be read in before
kanamycin_cgenff_toppar.str: kanamycin CHARMM format topology and parameters, requires cgenff parameters to be read in before
lymecycline_cgenff_toppar.str: lymecycline CHARMM format topology and parameters, requires cgenff parameters to be read in before
metacycline_cgenff_toppar.str: metacycline CHARMM format topology and parameters, requires cgenff parameters to be read in before
micronomicin_cgenff_toppar.str: micronomicin CHARMM format topology and parameters, requires cgenff parameters to be read in before
minocycline_cgenff_toppar.str: minocycline CHARMM format topology and parameters, requires cgenff parameters to be read in before
netilmicin_cgenff_toppar.str: netilmicin CHARMM format topology and parameters, requires cgenff parameters to be read in before
omadacycline_cgenff_toppar.str: omadacycline CHARMM format topology and parameters, requires cgenff parameters to be read in before
oxytetracycline_cgenff_toppar.str: oxytetracycline CHARMM format topology and parameters, requires cgenff parameters to be read in before
paromomycin_cgenff_toppar.str: paromomycin CHARMM format topology and parameters, requires cgenff parameters to be read in before
pipacycline_cgenff_toppar.str: pipacycline CHARMM format topology and parameters, requires cgenff parameters to be read in before
plazomicin_cgenff_toppar.str: plazomicin CHARMM format topology and parameters, requires cgenff parameters to be read in before
ribostamycin_cgenff_toppar.str: ribostamycin CHARMM format topology and parameters, requires cgenff parameters to be read in before
rolitetracycline_cgenff_toppar.str: rolitetracycline CHARMM format topology and parameters, requires cgenff parameters to be read in before
sarecycline_cgenff_toppar.str: sarecycline CHARMM format topology and parameters, requires cgenff parameters to be read in before
sisomicin_cgenff_toppar.str: sisomicin CHARMM format topology and parameters, requires cgenff parameters to be read in before
spectinomycin_cgenff_toppar.str: spectinomycin CHARMM format topology and parameters, requires cgenff parameters to be read in before
streptomycin_cgenff_toppar.str: streptomycin CHARMM format topology and parameters, requires cgenff parameters to be read in before
tetracycline_cgenff_toppar.str: tetracycline CHARMM format topology and parameters, requires cgenff parameters to be read in before
tigecycline_cgenff_toppar.str: tigecycline CHARMM format topology and parameters, requires cgenff parameters to be read in before
tobramycin_cgenff_toppar.str: tobramycin CHARMM format topology and parameters, requires cgenff parameters to be read in before
